# The field of protein function prediction as viewed by different domain scientists

**DOI:** 10.1101/2022.04.18.488641

**Authors:** Rashika Ramola, Iddo Friedberg, Predrag Radivojac

## Abstract

Experimental biologists, biocurators, and computational biologists all play a role in characterizing a protein’s function. The discovery of protein function in the laboratory by experimental scientists is the foundation of our knowledge about proteins. Experimental findings are compiled in knowledge-bases by biocurators to provide standardized, readily accessible, and computationally amenable information. Computational biologists train their methods using these data to predict protein function and guide subsequent experiments. To understand the state of affairs in this ecosystem, centered here around protein function prediction, we surveyed scientists from these three constituent communities. Our objective was to understand their views on this research area, including the importance of the problem, the usefulness of the methods, the bottlenecks in the field, and the level of interaction between the communities. We show that the three core communities have common but also idiosyncratic perspectives on the field. Most strikingly, experimentalists rarely use modern prediction software, but when presented with predictions, report many to be surprising and useful. Ontologies appear to be highly valued by biocurators, less so by experimentalists and computational biologists, yet controlled vocabularies bridge the communities and simplify the prediction task. Additionally, many software tools are not readily accessible and the predictions presented to the users can be broad and uninformative. To meet both the social and technical challenges in the field, a more productive and meaningful interaction between members of the core communities is necessary.

## 1 Introduction

A major objective of the field of computational biology is to develop theory, methodology, and software that support and drive biological discovery. The field has recently reported notable successes, including the development of tools for genome assembly (Nagarajan and Pop, 2013), methods for deriving the tree of life (Hinchliff et al., 2015), or deep-learning pipelines for accurately inferring a protein’s three-dimensional structure (Jumper et al., 2021). Although still expanding, one of the long-standing directions, yet still at the forefront of the field, is the development of techniques towards understanding the full repertoire of a protein’s activity under different molecular and environmental conditions (Shehu et al., 2016). Prominent examples of proteins with well-understood *function* include hemoglobin, an oxygen transporter (Antonini and Brunori, 1970), the tumor suppressor protein p53 that is altered in many cancers (Miyashita et al., 1994), or SARS-CoV-2 spike protein that enables the entry of the virus into mammalian cells (Hoffmann et al., 2020).

Protein function research is heavily invested in the study of model organisms. These organisms (e.g., yeast, mouse) are used for in-depth understanding of biological systems, with the expectation that discoveries made in a model organism will offer insights into the workings of other organisms. Although this strategy is reasonable, functional annotations accumulate slowly and remain far from complete even in model species. Filling this knowledge gap not only requires accurate prediction, but also systematization of knowledge (International Society for Biocuration, 2018), computational resource prioritization (Kacsoh et al., 2019) and statistical analysis (Subramanian et al., 2005). Computational researchers are therefore in a prominent position to build upon all experimental and curated data and develop inference systems for the understanding of protein function.

Historically, protein function has been described in the literature using natural language, but to unify terminology across all domains of life and to standardize computational pipelines, major knowledgebases, such as UniProtKB (The UniProt Consortium, 2019), often report function using ontological representations (Robinson and Bauer, 2011). An ontology is typically formalized as a general-to-specific concept hierarchy represented by a directed acyclic graph, where nodes are associated with textual descriptors (concepts, terms) that are mutually connected by relational ties of different types (e.g., is-a, part-of). The computational biology community has adopted the Gene Ontology (GO) (Ashburner et al., 2000) to describe function of biological macromolecules and the computational protein function prediction can therefore be seen as the problem of annotating a protein sequence with a subset of GO terms (Figure 1).

**Figure 1:**
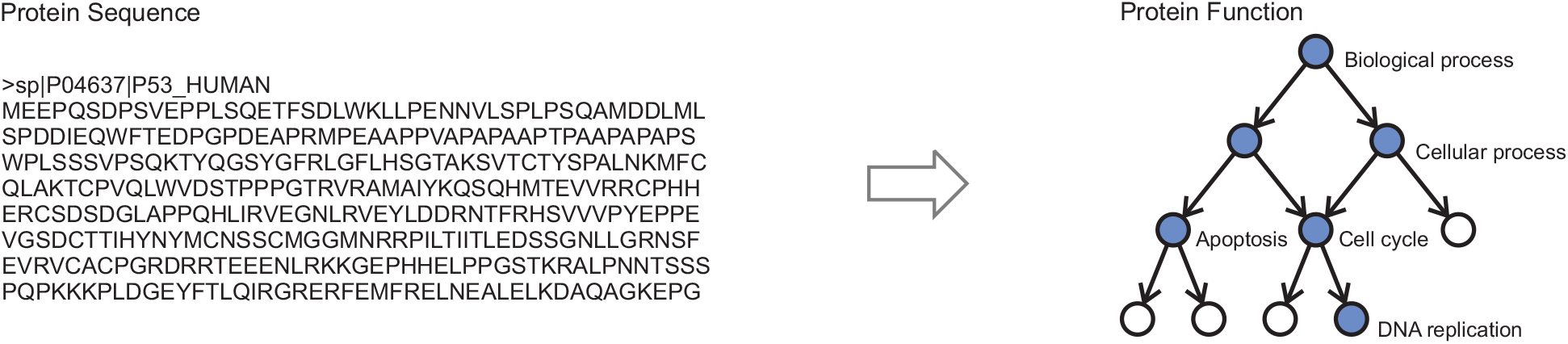
Stylized representation of the protein function prediction problem. (Left) Protein sequence of the human cellular tumor suppressor protein p53 consisting of 393 amino acids. The protein is shown in a FASTA format. (Right) Protein function defined as a set of blue nodes in the ontology graph consisting of 10 nodes. Each node is associated with a textual descriptor (term) from a controlled vocabulary; e.g., “biological process” as the root, “apoptosis” as an internal node, and “DNA replication” as a leaf node. The terms in the ontology are connected by relational ties such as is-a, that are read in the direction opposite the arrow. The task for a function prediction algorithm is to take an input sequence and output the nodes (terms) colored in blue. Note that the prediction of ontological terms is a somewhat simplified view of function prediction as it, for the most part, ignores the contextual, spatio-temporal and causal relations.

There are three core domain-expertise communities that are associated with protein function prediction. The first community is experimental biologists, who determine the function of proteins using experimental assays (Weber, 2004). The second is biocurators, who develop ontologies to describe protein function and generate data repositories of annotated proteins based on experimental and computational evidence (International Society for Biocuration, 2018). The third is computational biologists, who develop algorithms and software to predict a protein’s function from its amino acid sequence, structure, gene expression, genomic context, scientific literature, and more Radivojac et al. (2013). In some cases, members of these communities overlap, such as a lab that has an experimental component and also develops algorithms, or biocurators who are involved in software development. But in most cases members of each community specialize in one of these three domains.

In this work, we set out to understand how members of each of the three communities work on understanding protein function, how they perceive computational function prediction, how well they understand and use the work produced by members of the other communities, and how much they interact with each other. To do so we surveyed members of each of these communities using general as well as individualized questionnaires. Here we summarize major findings from these surveys that we believe describe the state of affairs in the field. We argue that a deeper understanding of how this scientific field operates, and where the bottlenecks may be, will lead to an improved quality of interaction between three constituent communities and ultimately advance the technical component of the field itself.

## 2 Methodology

The assessment of the field of protein function prediction was carried out using five online questionnaires—three for experimental scientists and one each for biocurators and computational biologists. The consent forms and the surveys were presented to the participants using Qualtrics (Supplementary Materials), and each person was offered a $10 gift card for participation. The study protocol was approved by the Institutional Review Board of Northeastern University (IRB# 19-10-08).

### 2.1 Participants

We reasoned that only protein function experts have deep insight into the field and therefore targeted active doctorate-level participants. We required interest and published work in the protein function domain, measured by the the reputation of the publishing venue. Participants were recruited via emails that briefly described the purpose of the study and laid out the expectations for the online questionnaires. The initial email was personalized and included a link to the consent form and subsequent survey materials. We also assured potential participants that their privacy would be protected.

We approached 169 scientists from 94 institutions using our understanding of the field as well as recommendations from colleagues with knowledge in the field (Table 1). Of the 169 individuals, 86 consented to take the survey, and 82 completed the survey (22 experimentalists, 19 biocurators, and 41 computational biologists). Multiple additional participants engaged in email exchanges with us, but indicated lack of time to complete the survey. The moderate number of overall responses is due to several reasons: (i) the survey was designed for specialists in the field; (ii) the individualized nature of parts of the questionnaires was designed to apply to the specialty of the respondents, which could not be done *en masse*; (iii) completing the questionnaires in full required a commitment not many respondents could provide.

**Table 1:**
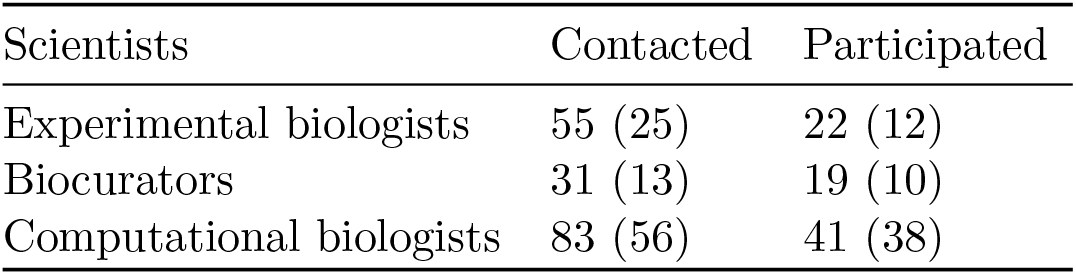
Basic statistics on the number of participants in the study. The number of institutions with which the participants are affiliated is parenthesized.

### 2.2 Surveys for experimental scientists

The three questionnaires for the experimental scientists are referred to here as the “general”, “specific”, and “comparative” questionnaires. The general questionnaire inquired about the background information and overall attitude towards protein function annotation and prediction (Section 2.2.1). The specific questionnaire was designed to assess the quality of the state-of-the-art function predictions on the proteins that are experimentally studied by consented domain experts; therefore, it was tailored to each surveyed participant (Section 2.2.2). We examined each person’s publications for proteins they investigated and asked each scientist to provide two proteins in which they believe they have the most expertise; and these proteins were subsequently used in the questionnaire. Finally, the comparative questionnaire was designed to assess the quality of the predictions of state-of-the-art methods relative to the predictions from baseline methods in the field (Section 2.2.3). The comparative questionnaire was also individualized.

#### 2.2.1 General Questionnaire

The general questionnaire was designed to understand the research background of the subjects, the level of their familiarity with bioinformatics software and databases, their understanding of GO annotations, and their need for protein function prediction. Overall, the questionnaire contained 28 multiple-choice questions and 10 free-text questions. Each participant was asked to answer the same set of questions. The median time of completion was 10 minutes.

#### 2.2.2 Specific Questionnaire

The specific questionnaire was developed to understand the performance of function prediction algorithms, as seen by experimentalists. This questionnaire was individualized with the rationale that only an expert can judge the quality, usefulness, and surprise of computational predictions. Each participant was first asked to self-assess their knowledge and then to assess the performance of one of the state-of-the-art algorithms on four proteins. Two of the proteins, *α*-hemoglobin and p53, for which we expected every participant to have rudimentary knowledge of their function, were identical for all participants. The remaining two proteins were unique to each participant and were selected from their own publications. We expected the researchers to have in-depth knowledge of the function of proteins they actively studied.

The state-of-the-art algorithm was selected randomly for each researcher from four tools that have been found to perform well (Radivojac et al., 2013; Jiang et al., 2016; Zhou et al., 2019), that represent methodological diversity in the field, and are regularly used. These included PFP (Hawkins et al., 2006), FFPred (Cozzetto et al., 2016), NetGO (You et al., 2019), and DeepGOPlus (Kulmanov and Hoehndorf, 2020). These methods leverage machine learning to learn protein function from a protein’s amino acid sequence, its 3D structure, or its network of interactions with other molecules. Due to their own strengths and limitations, we randomly picked one tool for each participant.

Researchers were presented with up to 25 highest-scoring terms for each of the proteins on each domain of GO (molecular function, biological process, cellular component). They were then asked to click on one or more of the following characterizations for each predicted term: (i) “known”, (ii) “useful”, (iii) “surprising, possible”, (iv) “surprising, doubtful”, and (v) “wrong”. We subsequently asked them to tell us if a predictor did a good job and then to use a text field to offer any additional information about the protein that was absent from the set of terms outputted by the predictor. We next asked them to describe the steps they typically take when they study function of a protein.

Overall, the specific questionnaire presented 10 questions to the researchers. Four questions were answered by clicking among pre-specified options and 10 were textual fields with free-form answers. However, as described above, the multiple choice questions contained a number of predicted terms (four proteins presented to the researcher, three GO domains), thus requiring significant time to complete. The median time to completion was 50 minutes.

#### 2.2.3 Comparative Questionnaire

The comparative questionnaire was developed to contrast the predictions from the state-of-the-art algorithm researchers saw in the specific questionnaire against the predictions from one of the baseline algorithms. We considered three different methods as baselines and randomly picked one for each researcher: (i) Naïve (Clark and Radivojac, 2011), (ii) TOPBLAST (Martin et al., 2004), and (iii) GOtcha (Martin et al., 2004). The predictions from the state-of-the-art tool and a baseline tool were presented side-by-side. The participants were then asked to evaluate the performance of the two tools relative to each other. Previous work has shown that, numerically, the advanced methods outperform baselines (Radivojac et al., 2013; Jiang et al., 2016; Zhou et al., 2019). Our objective here was to understand whether these numerical differences translate to meaningful information for domain experts.

Overall, the comparative questionnaire presented 37 questions to the researchers. Thirty-six questions were answered by clicking among pre-specified options and 1 was a textual field with free-form answers. The median time to completion of this survey was 6 minutes.

### 2.3 Surveys for biocurators and computational biologists

The questionnaires for biocurators and computational biologists were developed to understand the background of the participants, their familiarity with databases and software used in bioinformatics, their opinion of the GO terms and evidence codes, their level of interaction with other communities, the feedback that they have received about their research products, and their thoughts about the Critical Assessment of Function Annotation (CAFA). In contrast to experimentalists, biocurators and computational biologists were not required to enter their name in the survey and therefore could stay anonymous.

We asked both groups whether they think protein function prediction is an important problem and what they think are the bottlenecks in the field. Since both communities are related to CAFA (Friedberg and Radivojac, 2017), we also asked them about their familiarity with CAFA and their opinion of the evaluation metrics used in CAFA. We asked the computational biologists whether their lab developed tools for function prediction, and if they did, then what were the distinctive features of their algorithm, how they think the results of the protein function prediction pipeline should be presented and what steps they think experimentalists should follow when investigating a protein. Some of the questions asked to the biocurators were how they envision scientists using GO, whether they think experimental scientists use ontologies and databases the way they envision, and what they think are the typical misuses of GO.

An additional set of questions were asked of the biocurators about their reaction to the output of a protein function prediction algorithm (a list of GO terms and scores) to hemoglobin. This was to assess how they, as the developers of ontologies, would react to such usage of GO terms. The goal was to identify the differences in the intended usage and actual usage of ontologies as perceived by the biocurators. Biocurators were asked how useful they think these predictions are when used to convey protein function in that manner, how important it is for the bioinformatics community to build such algorithms, what should computational biologists keep in mind about ontologies when developing tools, how useful these predictions are for experimental biologists, and what experimental biologists should keep in mind when using such predictions.

The median time for the completion of the surveys was 46 minutes (38 questions) for the biocurators and 45 minutes (36 questions) for the computational biologists.

### 2.4 Statistical significance and confidence intervals

Asymmetric 68% confidence intervals in all evaluations were estimated using bootstrapping with 1,000 iterations of the entire cohort (Efron and Tibshirani, 1986). Statistical significance was determined using the *χ*^2^-test or t-test, as appropriate.

## 3 Results

### 3.1 Participants’ expertise and training

To learn about the participants we were working with, we asked about their years of research experience and the field(s) of expertise. As shown in Figure 2, the majority were highly experienced, and with expertise in several fields, but predominantly Biology and Computer Science.

**Figure 2:**
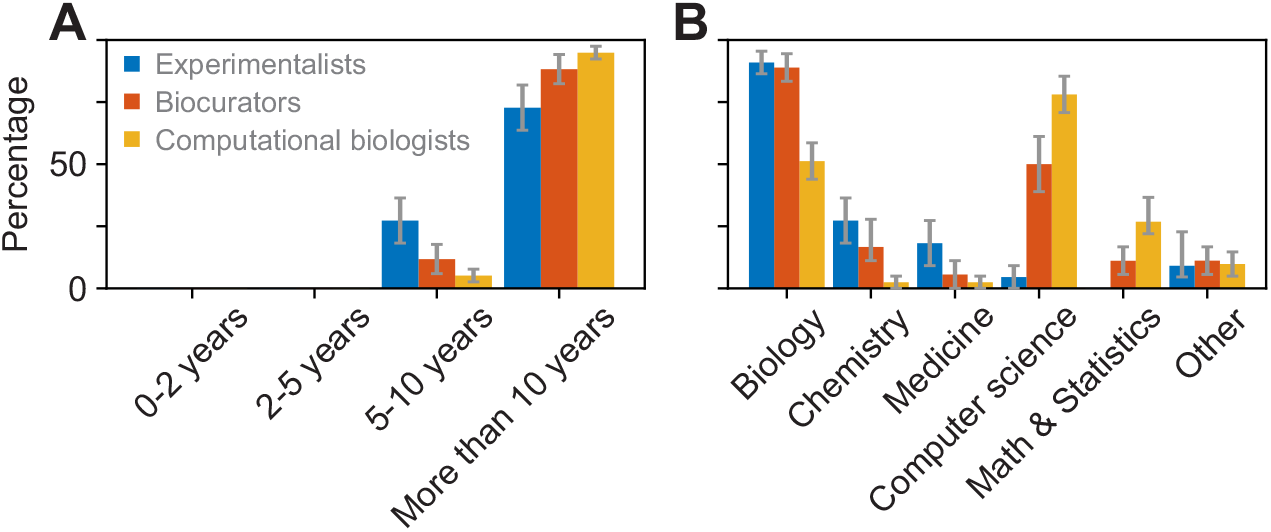
Experience and expertise of the study participants. Blue, red, and yellow bars show the answers from experimentalists, biocurators, and computational biologists, respectively. (A) Years of research experience and (B) Field(s) of training, where participants were allowed to select any number of training areas.

To establish type of expertise, we asked about the familiarity with common protein annotation databases and software. The participants were first offered a list of databases and asked to specify the level of familiarity with them, then they were asked to list additional databases they work with. Overall, biocurators appear to be more familiar with all databases, and especially with model-organism specific databases (Figure 3). A more equal, and high level of familiarity was shown for the general databases such as UniProtKB, Swiss-Prot, GO, and PDB (Berman et al., 2000). The more specialized model organism, enzyme, and metagenomic databases were less familiar to experimentlists. This is likely because model organism databases are only familiar to the experimentalists working with those model organisms, whereas biocurators are more familiar with all of them, as this knowledge is inherent to the biocuration community. The same applies to the more specific domain-oriented databases.

**Figure 3:**
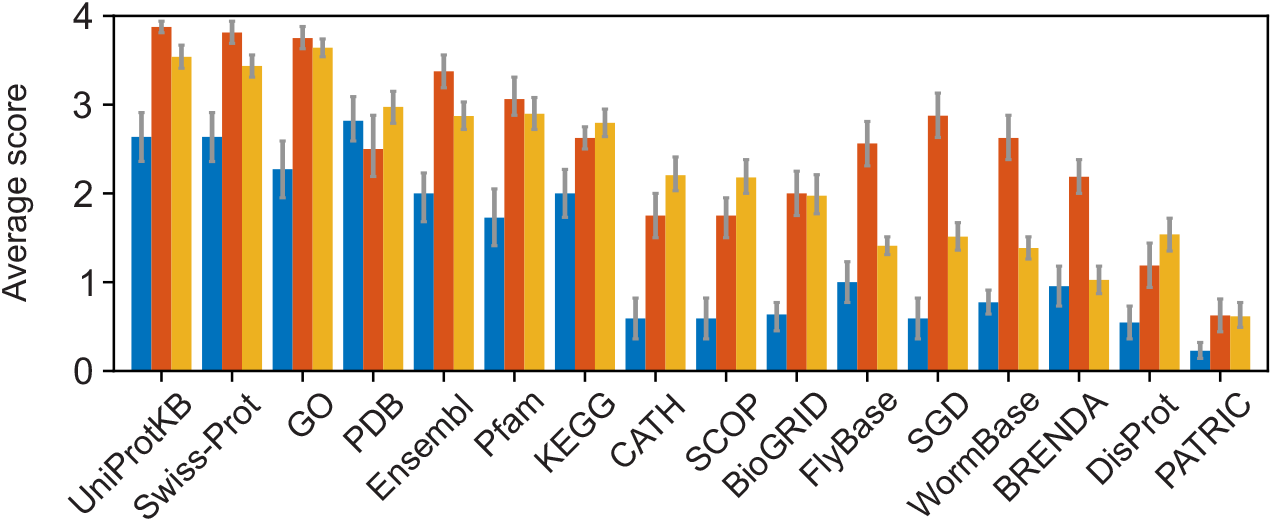
Familiarity with major databases. Each bar represents an average score for a core group indicating the familiarity with databases, where 0 = “not familiar”, 1 = “heard of it”; 2 = “use rarely”; 3 = “use sometimes”; 4 = “use frequently”. Blue, red and yellow bars show the average scores from experimentalists, biocurators and computational biologists, respectively.

In terms of bioinformatics software, we found that the participants from all three communities use numerous software packages. Of the tools related to function prediction, all were equally versed with BLAST (Altschul et al., 1997), with an average score of 3.6 ± 0.3, 3.4 ± 0.4, and 3.5 ± 0.3, by experimentalists, biocurators, and computational biologists, respectively. As with BLAST, no significant difference in familiarity with ClustalW (Thompson et al., 1994) and similar packages was found. The textual answers list tools used by the researchers, though the use of software was unequal. Asked to discuss what bioinformatics software is used by the researcher, one experimentalist responded with “Too many to list - mostly for NGS analysis […]”, while another with “None– I ask a colleague or get someone in my lab to ask them.”.

### 3.2 Experimentalists rarely use new prediction software

To gauge familiarity of experimental scientists with the use of protein function prediction software, we asked “Do you use any software packages for the purpose of understanding protein function?” Only 8/22 (36%) participants responded with a “yes”, whereas 14/22 (64%) responded with a “no” (Figure 4). This is a surprising outcome given the existence of over a hundred available methods developed by the bioinformatics and machine learning communities, of which dozens are readily available using web interfaces and others as downloadable software. In contrast, a deeper investigation revealed that all 22/22 (100%) researchers were familiar with BLAST, a tool featured in two of the baseline algorithms for function prediction (Section 2).

**Figure 4:**
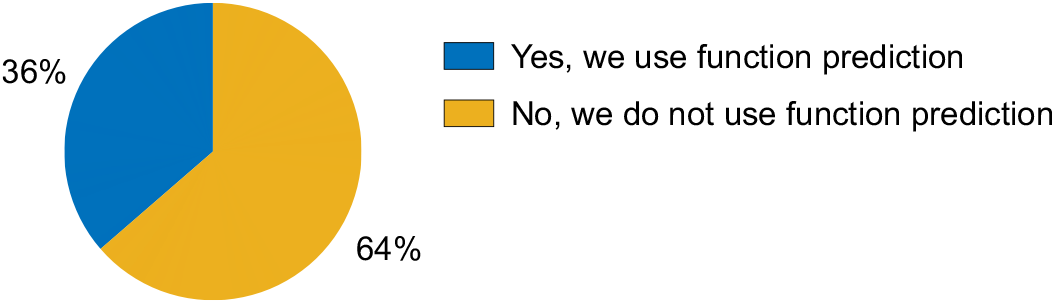
Use of function prediction software by experimentalists. The pie chart summarizes answers to the question “Do you use any software packages for the purpose of understanding protein function?”

The researchers who answered with a “yes” for the use of function prediction tools were further asked to list the tools that they use for this purpose. The answers included PSI-BLAST (Altschul et al., 1997), InterPro (Blum et al., 2021), HHpred (Zimmermann et al., 2018), STRING (Szklarczyk et al., 2021), Swiss-Prot (The UniProt Consortium, 2019), and I-TASSER (Roy et al., 2010). Of these, PSI-BLAST, InterPro via InterProScan, HHpred, and I-TASSER can be considered tools that help understanding of protein function, whereas STRING and Swiss-Prot are databases that provide computational predictions in a manner similar to experimental annotations.

Overall, we find that a small fraction of experimental researchers actually use advanced machine learning tools to understand protein function. Along the same lines, we identified some misunderstanding among experimental researchers as to what protein function prediction tools are.

### 3.3 Many predictions found useful by experimentalists

Here we summarize the reactions of experimental researchers to the predictions of state-of-the-art tools. Each participant was given up to 25 of the highest-scoring predictions by one of the computational tools (Section 2). That is, for each of the four proteins (hemoglobin, p53 and two they study experimentally in their own lab) they were shown up to 25 predictions for each of the three GO domains (molecular function, biological process, cellular component). For each such predicted term we provided the numerical score given by the algorithm and offered the following check boxes: “known”, “useful”, “surprising, possible”, “surprising, doubtful” and “wrong”, where a selection of multiple boxes or no boxes at all was allowed.

The summary of respondents’ assessments of individual predictions is shown in Figure 5A. It is not surprising that “known” was the predominant answer (2,049 terms) because some of the proteins were well-researched and the predictors may have been predicting on their own training data as well. More interestingly, 705 terms were found to be useful and 645 to be surprising and possible. This is an important finding that suggests the ability of the advanced methods to offer actionable information to a biomedical researcher, of which some predictions would be unexpected. Another 653 and 765 terms were found to be doubtful or incorrect, suggesting that there is considerable room to improve the methods and that the predictions that would be useful for domain experts are often mixed with a similar number of incorrect predictions.

**Figure 5:**
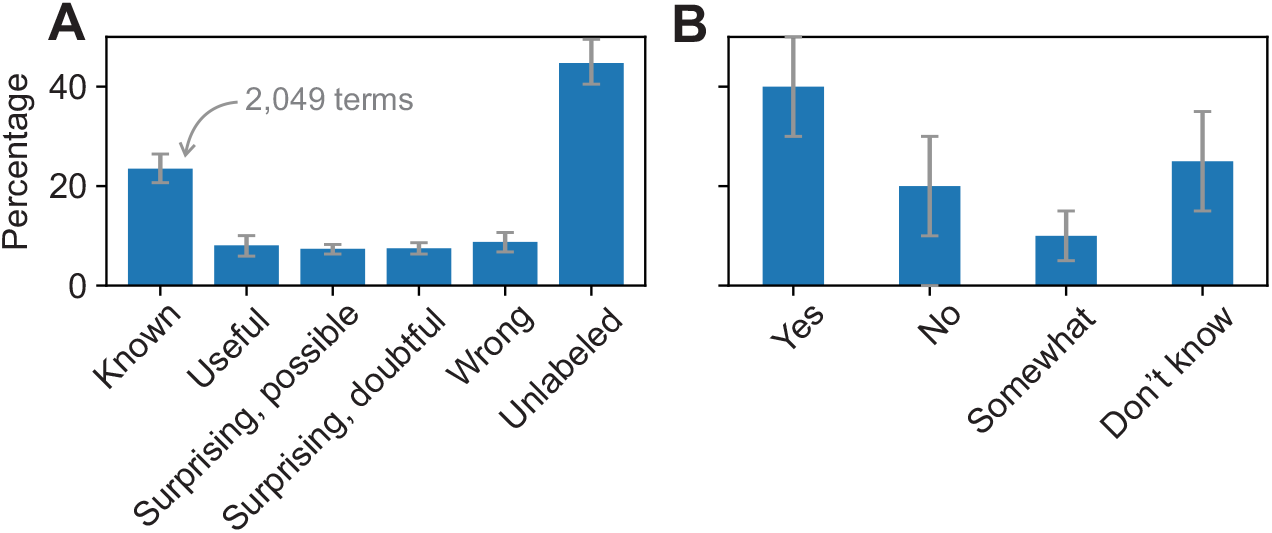
Assessment of the quality of predictions of the state-of-the-art software. (A) Percentage of terms from all three domains of GO presented to experimentalists they labeled by a particular descriptor. The “unlabeled” category shows the terms that have not been assigned any designation by the participants; e.g., skipped. Roughly half of each category corresponds to well-researched proteins hemoglobin and p53, whereas others correspond to the two specific proteins that each individual studies experimentally. (B) Percentage of experimentalists characterizing the prediction of state-of-the-art protein function predictors in response to the survey question “Do you think the algorithm has done a good job?”.

The experimental scientists were then asked to assess if the software has done a good job. A plurality (40%) of researchers responded that the algorithm has in fact done a good job with an additional 10% answering that the algorithm has done a somewhat good job. About 20% responded “no”, whereas about 30% of participants decided to either not answer this question or answer it with uncertainty (Figure 5B).

The participants were also asked to write free-text answers to describe what they think about the predictions. A representative set of criticisms includes “Extremely superficial annotation - some is super-obvious and likely only useful in superficial classifications.”, “I don’t think any of this is really all that useful–most of what comes out is things like the protein binds a metal ion, well most proteins do that, its kinda like saying that they’re made of amino acids.”, “I found these descriptions redundant and unhelpful (but also promising) in many ways.”, and “This is an enzymatic domain (metallopeptidase) that has lost key catalytic residues is likely to be inactive. It regulates other enzymes instead.”. These answers highlight the difficulty of the protein function prediction task, the difficulty of presenting the results to a downstream user as well as fairly high expectations from experimental scientists. The high expectations are exemplified by the last of the quotes above where a protein that looks like an enzyme has in fact evolved to lose the catalytic ability by mutations of a few key amino acids. To our knowledge, such fine-grained prediction by the machine learning and other computational tools has not yet been achieved.

### 3.4 Experimentalists do not distinguish between state-of-the-art and baseline algorithms

Extensive evaluation of protein function prediction revealed small but a consistently better performance of the state-of-the-art algorithms compared to the baseline methods (Radivojac et al., 2013; Jiang et al., 2016; Zhou et al., 2019). While the metrics used for evaluation of algorithms are broadly considered reasonable, it is unclear to what extent an improved numerical performance translates into the same characterization by experimental scientists. In this part of the survey, each experimental scientist was shown side-by-side predictions from the same state-of-the-art algorithm they had previously seen and one of the baseline tools for the same protein. Then, they were asked to assess the performance of the two algorithms. These results are summarized in Figure 6.

**Figure 6:**
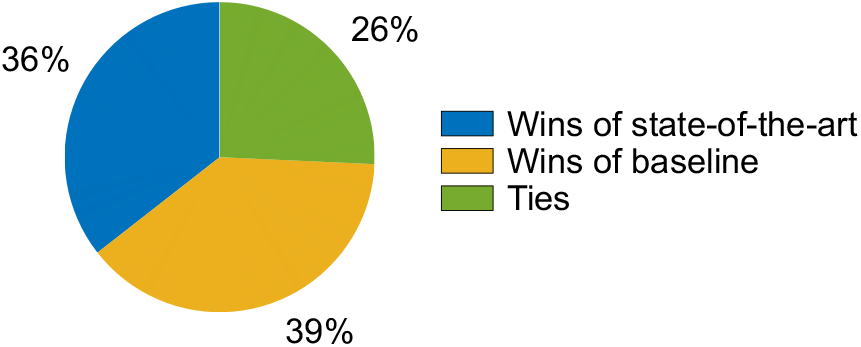
Summary of the comparisons between state-of-the-art and baseline tools. The pie chart shows a relatively equal number of wins between the state-of-the-art and the baseline algorithm. State-of-the-art algorithms were found to be better than Naive, but slightly worse than TOPBLAST and GOtcha.

These results suggest that, at this time, a numerical assessment of function prediction does not fully mirror an expert’s assessment. Although these results could be confounded (e.g., we showed only a list of top 25 predictions in each ontology of GO to reduce the burden of participation), this is a surprising finding that suggests some examination is needed of the metrics used for quality assessment of protein function prediction, the ontologies used to describe function, as well as the ways predictions are presented to experimental scientists.

### 3.5 Opinion of ontologies

The biocuration community has developed ontologies to unify protein function studies across species and to facilitate computation (Robinson and Bauer, 2011). At the same time, experimental scientists predominantly use natural language to learn about other proteins and to describe their findings. Since controlled vocabularies are an essential part of the prediction process, we investigated the familiarity of all core groups of scientists with GO.

We first asked “How useful do you think is a GO annotation for an experimental scientist?”. While most participants classified GO as at least “somewhat useful”, the answers varied between the core communities, with experimental scientists showing a preference for literature search in the process of knowledge acquisition and having a different trend than biocurators and computational biologists (Figure 7A). We next asked “How well do you think GO terms describe protein function?”. Most experimental and computational researchers answered “well enough”, and slightly differed between “not well at all” and “very well” (Figure 7B). Biocurators significantly differed in their opinion, most of whom considered that GO terms describe the function “very well”.

**Figure 7:**
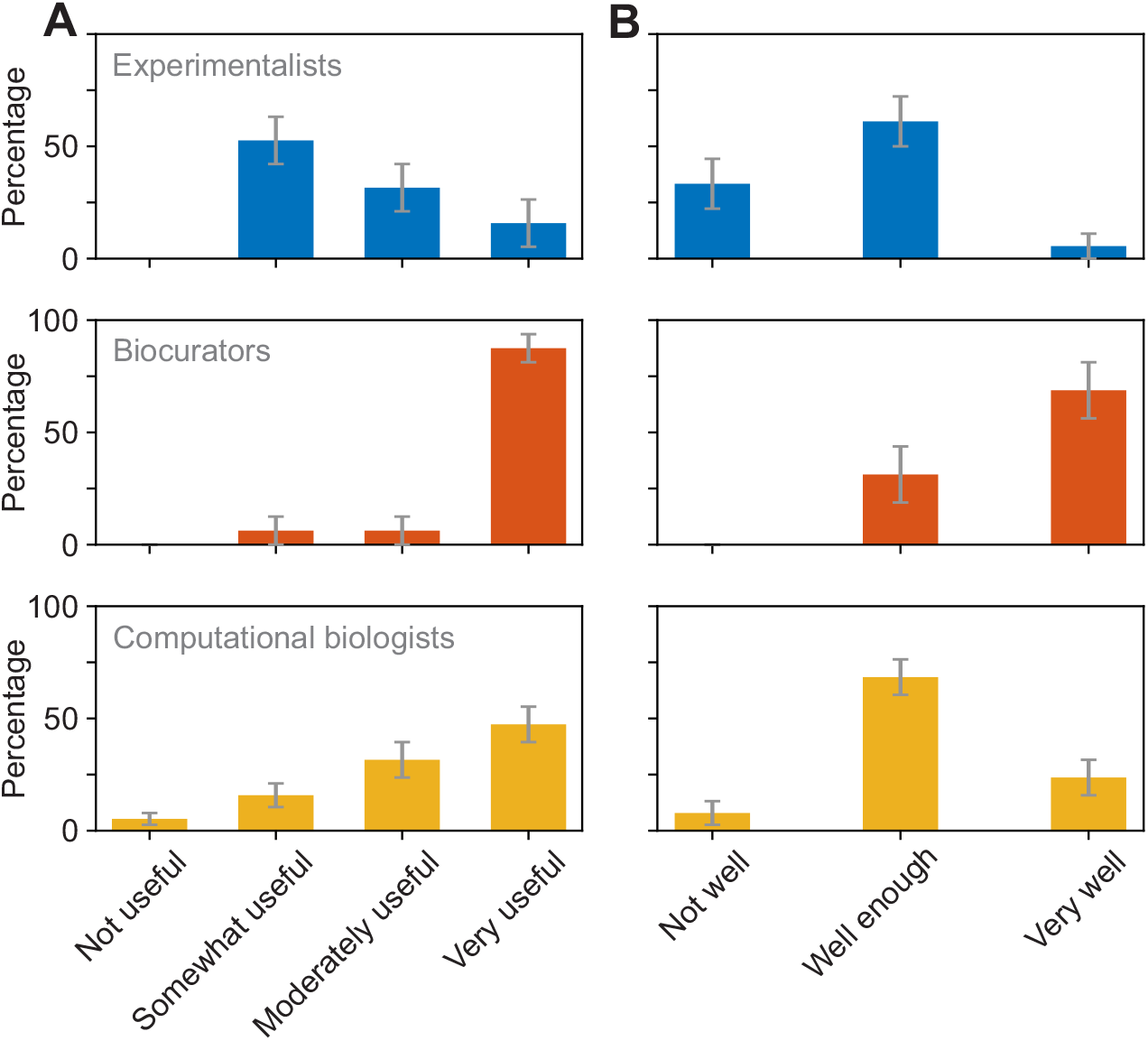
Perception of the utility of GO annotations. Blue, red, and yellow bars show the answers from experimentalists, biocurators, and computational biologists, respectively. (A) Summary of the answers to the question “How useful do you think is a GO annotation for an experimental scientist?”. The groups gave statistically different answers (*P* = 4.8 *·* 10^*−*4^; *χ*^2^-test). (B) Summary of the answers to the question “How well do you think GO terms describe protein function”. The groups gave statistically different answers (*P* = 7.9 *·* 10^*−*5^; *χ*^2^-test).

The free-text answers about protein function prediction further suggest that some experimentalists tied the outputs of a function prediction program to the perceived utility of GO itself. Some respondents see that as both a fault of the prediction algorithm and a fault of the GO term; e.g., experimental scientists noted “The resolution/granularity of GO annotation is too low to be useful” or “Terms are very broad. It would be useful to have more specific biological functions. I was surprised that nothing related to hypoxia came up for [protein]”. Another experimentalist noted “GO has become an insider joke for many in the field, likely however due to misuse of what the annotation tells us. I.e. use of root terms such as ‘biological process’ has been ridiculed repeatedly, but likely due to unclear use of GO. Has become too standard to just add to show some sort of analysis of a used dataset, but needs much more consideration in standard use (a fact that lots of people don’t seem to be aware of).”

General questions regarding function prediction were also asked of biocurators and computational biologists in their surveys. Supporting the views from experimentalists, one of the biocurators noted about their interaction with experimentalists “They are frustrated by the vast number of terms returned with a enrichment analysis - often more than the number of genes in the query list”. One computational biologist noted “Function is a loose term, therefore can only be predicted loosely.”. This exemplifies numerous problems in the field and relates to the bottlenecks (Section 3.6).

### 3.6 Importance of prediction and bottlenecks in the field

Our next question was designed to examine the motivation of function prediction among biocurators and computational biologists, who best understand the function prediction field. We asked “Do you think that developing tools for protein function prediction is an important problem?” and allowed the respondents to select answers from “not important” to “key to driving biology”. Both groups answered the question with effectively the same distribution, with about 60% of them giving the highest importance and 30% saying “quite important”.

We next asked “What do you think are the chief bottlenecks in protein function prediction?”, where we allowed each participant to give multiple answers or no answers at all. We also provided a free-text field for those who selected “other”. We find that more biocurators than computational biologists believe that “methodology” and “evaluation” are the chief bottlenecks. On the other hand, more computational biologists than biocurators believe that “available data” and “ontologies” are the chief bottlenecks (Figure 8A). The latter distinction was statistically significant and coincides with the view that biomedical ontologies may be overly complex for the development of predictive algorithms (Peng et al., 2018).

**Figure 8:**
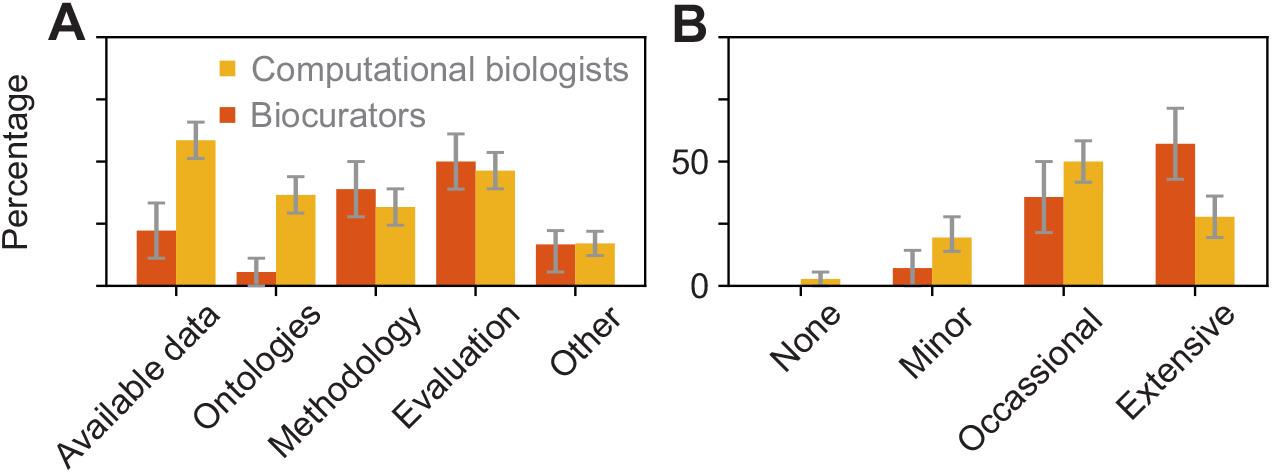
Bottlenecks in the function prediction field and levels of interaction between communities. Red and yellow bars show the answers from biocurators and computational biologists, respectively. (A) Summary of the answers to the question “What do you think are the chief bottlenecks in protein function prediction”. Statistically significant differences between groups were detected for “available data” (*P* = 2.2 *·* 10^*−*2^; t-test) and “ontologies” (*P* = 3.2 *·* 10^*−*2^; t-test). Categories were considered separately because each respondent could select any number of answers. (B) Summary of the answers to the question “How much do you interact with experimental scientists in your work”.

Among the researchers who selected “other”, the following answers stood out: “I would like to see progress in explainability – why did an algorithm come to its prediction. With some explainability, functional predictions would be better accepted by experimentalists.”, “Awareness of the area, other than in general terms. Use of the tools already developed in the area. Incorporation of predictions generated by these tools into standard databases like GO and HPO.”, “Lack of negative experimental examples […]”, and “Protein Identifier Cross-Referencing”.

Overall, we conclude that all factors have been often enough identified as the chief bottlenecks, including factors that extend beyond technical aspects of the field. Therefore, tangibly advancing any number of these areas is likely to be significant.

### 3.7 Level of interaction among core communities

Functional annotation of proteins is highly interdisciplinary, and making progress requires concerted work of researchers with domain knowledge in different disciplines. We therefore surveyed the participants regarding the level of collaborative work they perform with members of other core communities (Figure 8B). The most extensive interaction was reported between biocurators and computational biologists and least between experimentalists and computational biologists.

Among the free-text answers on how they interact with others and handle feedback from the userbase, some biocurators reported “Extensive feedback and interactions. These include suggestions for enhancement to the tools we maintain, reports of problems, requests for help which sometimes indicates we are missing a tools that would be useful to provide to our users.”, “They need extra terms added and we do so. We also get feedback on the submission process and we do our best to include the suggestions depending on resources and whether there is wider demand in the community for that new feature”, “A lot of the random requests about file formats/URLs are not addressable as we don’t have the budget to pull someone away from a central project to work on the request […] or translating the ontology to another language (especially non-latin alphabets).”, and “Our group […] has held roughly 50 onsite workshops with experts to build our ontology and have had over 7000 requests on our github tracker, so interaction has been pretty intense.”.

Computational biologists described their interaction as “We receive continuous feedback from users regarding many facets of our software (e.g., documentation, user interface, bug reports, etc.) We do our best to track and address user feedback using standard GitHub tools such as issues, pull requests and automated tests enabled by GitHub Actions.”, and “In one project, the agrobiotech company would run large-scale data-generating and validation experiments co-designed with us, and one of their staff meets weekly with my group throughout the project period discussing both technical ideas, results and refinements. In another project, it was more like one week a month kind of interactions as we progressed through different components of the project.”.

### 3.8 Assessment of critical assessments

Over the past 10 years, we have been organizing the CAFA challenge (Friedberg and Radivojac, 2017). Briefly, CAFA is an experiment designed to provide a large-scale assessment of computational methods dedicated to predicting protein function, using a prospective approach. CAFA organizers provide a large number of protein sequences and the predictors then predict the function of these proteins by associating them with ontological terms (Figure 1). Several months after the prediction deadline, the number of annotated sequences increases in the databases, and the predictions can be evaluated against the newly accumulated terms that have received experimental support. CAFA has become a community standard for assessing the performance of function prediction algorithms, with about 100 participating laboratories worldwide. As many of the biocurator and computational biologists survey respondents may have been familiar with CAFA, we solicited their opinions of the challenge. Specifically, we asked them about the usefulness of CAFA to the field, about evaluation metrics used in the assessments, as well as general questions.

We first asked “To what extent do you think CAFA is useful?” to which 60% of biocurators and 80% of computational biologists answered “highly useful”, 35% and 15% answered “somewhat useful”, and 5% of computational biologists did not find CAFA useful at all. About 10% of participating biocurators have never heard of CAFA (Figure 9A). These results show that an overwhelming majority in both groups find this community challenge beneficial for the field. Following the debate about the measures for the accuracy of function prediction (Dessimoz et al., 2013; Jiang et al., 2014; Plyusnin et al., 2019), we also inquired about the quality of the assessment metrics. We asked “How good are evaluation metrics used in CAFA for protein function prediction?” and allowed for the following answers: “They do not capture anything relevant”, “They capture some relevant information”, “They capture enough information relevant for decision making”, and “They capture most relevant information”. The results show that most participants find assessment metrics to capture at least some relevant biological information, with computational biologists seeing the current metrics more favorable than the biocurators (Figure 9B).

**Figure 9:**
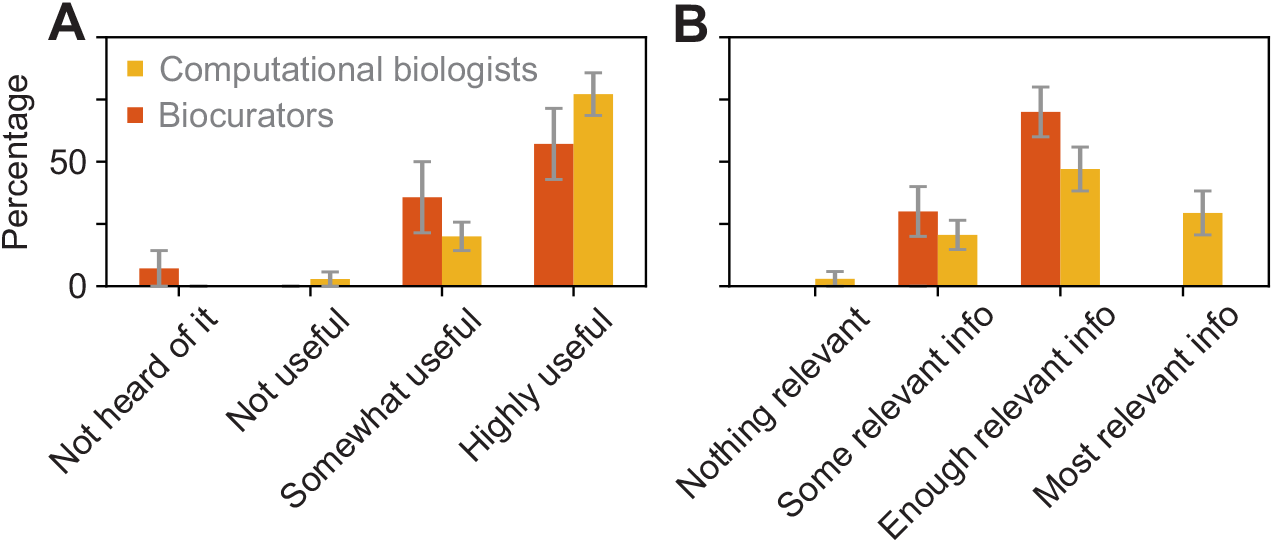
Assessment of CAFA. Red and yellow bars show the answers from biocurators and computational biologists, respectively. (A) Summary of the answers to the question “To what extent do you think CAFA is useful?”. (B) Summary of the answers to the question “How well do the evaluation metrics used in CAFA capture relevant information for protein function prediction?”. The differences between distributions are not significant.

Some free-text answers about CAFA include “CAFA has been extremely successful in driving this field - by ensuring methods are benchmarked properly and evaluated against each other. Also, in helping predictors test different strategies to see how well they perform. It has also raised the profile of this research field and helped explain the value to the wider biological community through some very high impact publications. By showing which strategies work well, it has helped secure funding for those initiatives.”, and “CAFA is indispensable to building a community and advancing the science. I look forward to more types of CAFA challenges toward defining and solving new problems related to protein function prediction (the recent addition of more ontologies was a good example). I also look forward to more CAFA-led community efforts to outreach, such as community-prediction of SARS-CoV-2 proteins when they were first sequenced last year or that of proteins found in meta genomes.”.

## 4 Related work

There is little literature assessing community experience in the field of bioinformatics. A 2019 investigation of 24,490 omics software packages revealed that more than a quarter were inaccessible via the original link provided in the publication (Mangul et al., 2019). Among a random sample of 99 of these tools, 49% were found “difficult to install” as they required more than 15 minutes of effort. The study reasoned that this is a direct consequence of the current research model in computational biology where publishing new tools and software is incentivized, whereas maintenance and production of tools that are easy to install and use is not. There have also been a few qualitative studies on usability of bioinformatics software and tools. Some were focused on the usability of a single resource such as a protein interaction database (Mirel, 2007) or a specific online resource that provides information on the evolutionary relationships of protein domains (Bolchini et al., 2009). These studies pointed out specific usability issues for these tools and offered recommendations for improvements in usability design. The lack and need of user-centered design in bioinformatics, where software is designed by taking the needs of the user into account has been pointed out by Pavelin et al. (2012).

While these previous studies point out usability issues in individual tools as well as the collective body of tools produced by the bioinformatics community, we are not aware of any research about the perception, characterization and use of protein function prediction by experimentalists. A study closest to ours was a qualitative survey of 11 life scientists that investigated their needs for bioinformatics software, but it did not investigate the perception of the quality of the methods (Morrison-Smith et al., 2015). Our study, at least with respect to experimentalists, has been tailored to the research questions asked of each participating scientist, so as to elicit expert opinion on the quality of predictions and potential use of protein function prediction in future research. It also surveyed all core communities involved in protein function prediction to understand the field more broadly.

## 5 Discussion

We present key findings from a study of three core communities involved in research of protein function: experimentalists, who generate new knowledge in the field by directly testing hypotheses; biocurators, who codify and standardise functional annotations; and computational biologists, who develop algorithms to predict function. To characterize a protein’s function in a way that is informative and useful, members of all three communities need to work together with an understanding of the type of work and the importance of their peers in other communities. In this work we found that while members of the three core domains have produced work that has advanced our understanding of protein function, there are a few areas in which collaboration can be improved.

One interesting finding was that experimentalists rarely use state-of-the-art machine learning software to predict function. While we did not survey for the reasons, it is reasonable to assume that several underlying causes are at play. First, the knowledge that such tools exist. While they are published in scientific literature, many of these tools do not gain visibility unless adopted by large go-to platforms such as those provided by NCBI or EBI. Second, the usability of prediction software. Software developed in research labs often lingers in the form of prototypes, tends to be cumbersome to use and is often not maintained regularly. Third, the question of need. While new machine learning software may have higher precision and recall than, say, BLAST or HHpred, BLAST and HHpred are often perceived as sufficient for the task at hand. Fourth, the trust and understanding. Even if experimentalists have access to new machine learning software, they still tend to trust highly cited baseline methods rather than a recently published method. Despite the reluctance and lack of time to find and apply advanced software, many predictions from such algorithms were found by experimentalists to be surprising and useful.

The survey of engagement between the domain communities offers some more information. Clearly, there is engagement between members of all communities. Computational biologists and experimentalists seem to have less interaction than any of these two groups with biocurators. Again, the reasons for this discrepancy are unclear, and may include that the inherent nature of biocuration requires work with experimentalists to properly capture function annotation and at the same time with computational biologists to provide data formatted in a computationally amenable manner. In contrast, computational biologists seem not to interact as much with experimentalists, which may be related to the fact that algorithmic advances are often more valued than software production in their home departments. Another reason is the lack of understanding of the needs of experimental communities and the different definitions of function used by experimentalists. Finally, the structure of academia is such that many interactions occur within a departmental level, and with like-minded peers, as opposed to interdisciplinary colleagues. These structural disincentives may also contribute to the lack of more productive and more meaningful interaction.

There are limitations to this work that include a relatively small selection of participants. Most of the answers provided by the consented participants were opinions and also incorporate their own interpretation of the intent of the question. We have found minor issues with the surveys in that some questions have in fact been misunderstood. Finally, despite our intent for otherwise, many participants took significant time to complete this work. The length of the survey may have confounded some of the findings; e.g., the comparative questionnaire for the experimentalists has taken much shorter time than other surveys. As the last of the three components, it is conceivable that some participants have lost focus by the end of the process.

That said, through this work we have discovered interesting trends in the interdisciplinary work within and between the three communities involved in functional annotation of proteins. While each community is producing admirable results, in the interest of accurate, reliable and sustained functional annotation, the quality of mutual interaction can be improved. Specifically, the state-of-the-art algorithms should be made visible and usable for experimentalists, which would stimulate more dialogue between experimental and computational groups. This would require adjustments in the way many computational groups are operating, and is best incentivized by funding agencies. Also, educating computational groups on the typical needs of the experimental communities, and conversely, experimentalists with basic computational skills would help to remove silos between the communities. The over- and mis-interpretation of ontologies also seems to be a problem, and the biocuration community is best positioned to address that.

To further advance the studies of protein function, and by extension understand the molecular underpinnings of life, we deem it important that both technical and social challenges of the field must be overcome.

## Acknowledgements

We thank Charles E. Dann, Justin D. Delano, Casey S. Greene and Sean D. Mooney for valuable help and insights in developing and executing this study.

## Notes

### Competing Interest Statement

The authors have declared no competing interest.

