## Supplementary Material for "The field of protein function prediction as viewed by different domain scientists"

### Supplementary Information

#### 1 The questionnaires for the experimental scientists

Here are the forms that were given to the experimental scientists:

- Consent Forms (Appendix A, E and G) - Since the questionnaires for the experimental scientists were tailored for each scientist individually, they were sent the consent form first. We made their individualised forms only after they consented to participate in this study.
- General (Appendix B) - This questionnaire was common for all the experimentalists. It had questions about their background, familiarity with bioinformatics databases and software, and their general perception of the gene ontology (GO) terms.
- Specific Form (Appendix C) - This questionnaire was engineered for each scientist based on the proteins of their expertise. The participants were provided with function predictions of four proteins - haemoglobin (chain alpha), P53 and two proteins of their expertise. We ask them to classify each protein into one or more of these five categories: (i) “known”, (ii) “useful”, (iii) “surprising, possible”, (iv) “surprising, doubtful”, and (v) “wrong”. The goal was to assess how they would react to each prediction term produced by a state-of-the art algorithm. Additionally, they were asked questions about the protein function prediction task and how they approach it. In the questionnaire linked here, the specific proteins have not been named to protect the privacy of the participants. They are referred to as ”Specific Protein 1” and ”Specific Protein 2”.
- Comparative Form (Appendix D) - This form was designed to assess how the predictions from the state-of-the art method would be perceived when compared to the predictions from a baseline method. Only the specific proteins (”Specific Protein 1” and ”Specific Protein 2”) were used in this form.

#### 2 Questionnaires for the biocurators and the computational biologists

Please see the questionnaire for biocurators (Appendix H), and the questionnaire for the computational biologists (Appendix F). The consent forms are included at the beginning of these surveys. These surveys are designed to ask the participants about their background, familiarity with bioinformatics databases and software, their perception of the Gene Ontology terms, how much they interact with other communities and their views about CAFA. In addition to these common questions, each community also received some tailored questions.

| S. No. | Database | URL (s) |
|---|---|---|
| 1 | UniProtKB [1] | https://www.uniprot.org |
| 2 | Swiss-Prot [2] | https://www.expasy.org/resources/uniprotkb-swiss-prot |
| 3 | GO [3, 4] | http://geneontology.org |
| 4 | PDB [5, 6] | http://www.wwpdb.org<br>https://www.rcsb.org |
| 5 | Ensembl [7] | https://useast.ensembl.org/index.html |
| 6 | Pfam [8] | http://pfam.xfam.org |
| 7 | KEGG [9] | https://www.genome.jp/kegg/pathway.html |
| 8 | CATH [10, 11] | https://www.cathdb.info |
| 9 | SCOP [12, 13] | https://scop.mrc-lmb.cam.ac.uk |
| 10 | BioGRID [14] | https://thebiogrid.org |
| 11 | FlyBase [15, 16] | https://flybase.org |
| 12 | SGD [17] | https://www.yeastgenome.org |
| 13 | WormBase [18] | https://wormbase.org/ |
| 14 | BRENDA [19] | https://www.brenda-enzymes.org |
| 15 | DisProt [20] | https://disprot.org |
| 16 | PATRIC [21] | https://www.patricbrc.org |

Table 1: The list of software which were displayed to the participants when asked ”Here is a list of some databases used in bioinformatics. Please indicate your level of familiarity with each one.”

| S. No. | Software | URL |
|---|---|---|
| 1 | BLAST [22] | https://blast.ncbi.nlm.nih.gov/Blast.cgi |
| 2 | CLUSTAL [23] | http://www.clustal.org |
| 3 | UCSC Genome Browser [24, 25] | https://genome.ucsc.edu |
| 4 | MEGA [26] | https://www.megasoftware.net |
| 5 | DNAStar | https://www.dnastar.com |

Table 2: The list of software which were displayed to the participants when asked ”Here is a list of some software packages used in bioinformatics. Please indicate your level of familiarity with each one.”

### 3 The Databases and Software listed in the surveys

To assess the familiarity of the participants with the field of bioinformatics, all the participants were about their familiarity with bioinformatics databases and software. They were asked to rate the databases (Table 1) and software (Table 2) as: 0 = “not familiar”, 1 = “heard of it”; 2 = “use rarely”; 3 = “use sometimes” or 4 = “use frequently”.

### Acknowledgements

We thank Charles E. Dann, Justin D. Delano, Casey S. Greene and Sean D. Mooney for valuable help and insights in developing and executing this study.

### References

[1] UniProt Consortium. UniProt: a worldwide hub of protein knowledge. *Nucleic Acids Res.*,

47(D1):D506–D515, January 2019.

[2] A Bairoch and R Apweiler. The SWISS-PROT protein sequence database and its supplement TrEMBL in 2000. *Nucleic Acids Res.*, 28(1):45–48, January 2000.

[3] The Gene Ontology Consortium. Gene ontology: tool for the unification of biology. *Nat. Genet.*, 25(1):25–29, May 2000.

[4] The Gene Ontology Consortium. The gene ontology resource: enriching a GOld mine. *Nucleic Acids Res.*, 49(D1):D325–D334, January 2021.

[5] Helen Berman, Kim Henrick, and Haruki Nakamura. Announcing the worldwide protein data bank. *Nat. Struct. Biol.*, 10(12):980, December 2003.

[6] Stephen K Burley, Helen M Berman, Gerard J Kleywegt, John L Markley, Haruki Nakamura, and Sameer Velankar. Protein data bank (PDB): The single global macromolecular structure archive. *Methods Mol. Biol.*, 1607:627–641, 2017.

[7] Kevin L Howe, Premanand Achuthan, James Allen, Jamie Allen, Jorge Alvarez-Jarreta, M Ridwan Amode, Irina M Armean, Andrey G Azov, Ruth Bennett, Jyothish Bhai, Konstantinos Billis, Sanjay Boddu, Mehrnaz Charkhchi, Carla Cummins, Luca Da Rin Fioretto, Claire Davidson, Kamalkumar Dodiya, Bilal El Houdaigui, Reham Fatima, Astrid Gall, Carlos Garcia Giron, Tiago Grego, Cristina Guijarro-Clarke, Leanne Haggerty, Anmol Hemrom, Thibaut Hourlier, Osagie G Izuogu, Thomas Juettemann, Vinay Kaikala, Mike Kay, Ilias Lavidas, Tuan Le, Diana Lemos, Jose Gonzalez Martinez, José Carlos Marugán, Thomas Maurel, Aoife C McMahon, Shamika Mohanan, Benjamin Moore, Matthieu Muffato, Denye N Oheh, Dimitrios Paraschas, Anne Parker, Andrew Parton, Irina Prosovetskaia, Manoj P Sakthivel, Ahamed I Abdul Salam, Bianca M Schmitt, Helen Schuilenburg, Dan Sheppard, Emily Steed, Michal Szpak, Marek Szuba, Kieron Taylor, Anja Thormann, Glen Threadgold, Brandon Walts, Andrea Winterbottom, Marc Chakiachvili, Ameya Chaubal, Nishadi De Silva, Bethany Flint, Adam Frankish, Sarah E Hunt, Garth R IIsley, Nick Langridge, Jane E Loveland, Fergal J Martin, Jonathan M Mudge, Joanella Morales, Emily Perry, Magali Ruffier, John Tate, David Thybert, Stephen J Trevanion, Fiona Cunningham, Andrew D Yates, Daniel R Zerbino, and Paul Flicek. Ensembl 2021. 49(D1):D884–D891, January 2021.

[8] Jaina Mistry, Sara Chuguransky, Lowri Williams, Matloob Qureshi, Gustavo A Salazar, Erik L L Sonnhammer, Silvio C E Tosatto, Lisanna Paladin, Shriya Raj, Lorna J Richardson, Robert D Finn, and Alex Bateman. Pfam: The protein families database in 2021. *Nucleic Acids Res.*, 49(D1):D412–D419, January 2021.

[9] Minoru Kanehisa. KEGG bioinformatics resource for plant genomics and metabolomics. *Methods Mol. Biol.*, 1374:55–70, 2016.

[10] Ian Sillitoe, Nicola Bordin, Natalie Dawson, Vaishali P Waman, Paul Ashford, Harry M Scholes, Camilla S M Pang, Laurel Woodridge, Clemens Rauer, Neeladri Sen, Mahnaz Abbasian, Sean Le Cornu, Su Datt Lam, Karel Berka, Ivana Hutařová Varekova, Radka Svobodova, Jon Lees, and Christine A Orengo. CATH: increased structural coverage of functional space. *Nucleic Acids Res.*, 49(D1):D266–D273, January 2021.

[11] Tony E Lewis, Ian Sillitoe, Natalie Dawson, Su Datt Lam, Tristan Clarke, David Lee, Christine Orengo, and Jonathan Lees. Gene3D: Extensive prediction of globular domains in proteins. *Nucleic Acids Res.*, 46(D1):D435–D439, January 2018.

[12] Antonina Andreeva, Dave Howorth, Cyrus Chothia, Eugene Kulesha, and Alexey G Murzin. SCOP2 prototype: a new approach to protein structure mining. *Nucleic Acids Res.*, 42(Database issue):D310–4, January 2014.

[13] Antonina Andreeva, Eugene Kulesha, Julian Gough, and Alexey G Murzin. The SCOP database in 2020: expanded classification of representative family and superfamily domains of known protein structures. *Nucleic Acids Res.*, 48(D1):D376–D382, January 2020.

[14] Rose Oughtred, Jennifer Rust, Christie Chang, Bobby-Joe Breitkreutz, Chris Stark, Andrew Willems, Lorrie Boucher, Genie Leung, Nadine Kolas, Frederick Zhang, Sonam Dolma, Jasmin Coulombe-Huntington, Andrew Chatr-Aryamontri, Kara Dolinski, and Mike Tyers. The BioGRID database: A comprehensive biomedical resource of curated protein, genetic, and chemical interactions. *Protein Sci.*, 30(1):187–200, January 2021.

[15] Jim Thurmond, Joshua L Goodman, Victor B Strelets, Helen Attrill, L Sian Gramates, Steven J Marygold, Beverley B Matthews, Gillian Millburn, Giulia Antonazzo, Vitor Trovisco, Thomas C Kaufman, Brian R Calvi, and FlyBase Consortium. FlyBase 2.0: the next generation. *Nucleic Acids Res.*, 47(D1):D759–D765, January 2019.

[16] J M McWhorter, E Alexander, C H Davis, and L Kelly. Posterior cervical fusion in chidren. *J. Neurosurg.*, 45(2):211–215, August 1976.

[17] J M Cherry, C Adler, C Ball, S A Chervitz, S S Dwight, E T Hester, Y Jia, G Juvik, T Roe, M Schroeder, S Weng, and D Botstein. SGD: Saccharomyces genome database. *Nucleic Acids Res.*, 26(1):73–79, January 1998.

[18] Todd W Harris, Igor Antoshechkin, Tamberlyn Bieri, Darin Blasiar, Juancarlos Chan, Wen J Chen, Norie De La Cruz, Paul Davis, Margaret Duesbury, Ruihua Fang, Jolene Fernandes, Michael Han, Ranjana Kishore, Raymond Lee, Hans-Michael Müller, Cecilia Nakamura, Philip Ozersky, Andrei Petcherski, Arun Rangarajan, Anthony Rogers, Gary Schindelman, Erich M Schwarz, Mary Ann Tuli, Kimberly Van Auken, Daniel Wang, Xiaodong Wang, Gary Williams, Karen Yook, Richard Durbin, Lincoln D Stein, John Spieth, and Paul W Sternberg. WormBase: a comprehensive resource for nematode research. *Nucleic Acids Res.*, 38(Database issue):D463–7, January 2010.

[19] Maurice Scheer, Andreas Grote, Antje Chang, Ida Schomburg, Cornelia Munaretto, Michael Rother, Carola Söhngen, Michael Stelzer, Juliane Thiele, and Dietmar Schomburg. BRENDA, the enzyme information system in 2011. *Nucleic Acids Res.*, 39(Database issue):D670–6, January 2011.

[20] Federica Quaglia, Bálint Mészáros, Edoardo Salladini, András Hatos, Rita Pancsa, Lucía B Chemes, Mátyás Pajkos, Tamas Lazar, Samuel Peña-Díaz, Jaime Santos, Veronika Ács, Nazanin Farahi, Erzsébet Fichó, Maria Cristina Aspromonte, Claudio Bassot, Anastasia Chasapi, Norman E Davey, Radoslav Davidović, Laszlo Dobson, Arne Elofsson, Gábor Erdős, Pascale Gaudet, Michelle Giglio, Juliana Glavina, Javier Iserte, Valentín Iglesias, Zsófia Kálmán, Matteo Lambrughi, Emanuela Leonardi, Sonia Longhi, Sandra Macedo-Ribeiro, Emiliano Maiani, Julia Marchetti, Cristina Marino-Buslje, Attila Mészáros, Alexander Miguel Monzon, Giovanni Minervini, Suvarna Nadendla, Juliet F Nilsson, Marian Novotný, Christos A Ouzounis, Nicolás Palopoli, Elena Papaleo, Pedro José Barbosa Pereira, Gabriele Pozzati, Vasilis J Promponas, Jordi Pujols, Alma Carolina Sanchez Rocha, Martin Salas, Luciana Rodriguez Sawicki, Eva Schad, Aditi Shenoy, Tamás Szaniszló, Konstantinos D Tsirigos, Nevena

Veljkovic, Gustavo Parisi, Salvador Ventura, Zsuzsanna Dosztányi, Peter Tompa, Silvio C E Tosatto, and Damiano Piovesan. DisProt in 2022: improved quality and accessibility of protein intrinsic disorder annotation. *Nucleic Acids Res.*, 50(D1):D480–D487, January 2022.

[21] James J Davis, Alice R Wattam, Ramy K Aziz, Thomas Brettin, Ralph Butler, Rory M Butler, Philippe Chlenski, Neal Conrad, Allan Dickerman, Emily M Dietrich, Joseph L Gabbard, Svetlana Gerdes, Andrew Guard, Ronald W Kenyon, Dustin Machi, Chunhong Mao, Dan Murphy-Olson, Marcus Nguyen, Eric K Nordberg, Gary J Olsen, Robert D Olson, Jamie C Overbeek, Ross Overbeek, Bruce Parrello, Gordon D Pusch, Maulik Shukla, Chris Thomas, Margo VanOeffelen, Veronika Vonstein, Andrew S Warren, Fangfang Xia, Dawen Xie, Hyunseung Yoo, and Rick Stevens. The PATRIC bioinformatics resource center: expanding data and analysis capabilities. *Nucleic Acids Res.*, 48(D1):D606–D612, January 2020.

### Appendices: Consent Forms and Questionnaires

Appendix A - Experimentalists: Consent Form 2

Appendix B - Experimentalists: General Form 4

Appendix C - Experimentalists: Specific Form 9

Appendix D - Experimentalists: Comparative Form 38

Appendix E - Computational Biologists: Consent Form 48

Appendix F - Computational Biologists: General Form 51

Appendix G - Biocurators: Consent Form 60

Appendix H - Biocurators: General Form 62

### Appendix A - Experimentalists: Consent Form

**Northeastern University, Khoury College of Computer Sciences**

**Name of Investigator(s):** Prof. Predrag Radivojac

**Title of Project:** Assessing the usability and value of protein function prediction algorithms

**Sponsor:** National Science Foundation

**Information Sheet**

**Request to Participate in Research**

We would like to invite you to participate in a web-based online survey. The survey is part of a research study whose purpose is to understand the perception and utility of computational protein function prediction methods for experimental scientists.

The surveys should take about 40 minutes to complete.

We are asking you to participate in this study because you are an experimental scientist with expertise in specific proteins. **You must be at least 18 years old to take this survey.**

**The decision to participate in this research project is voluntary.** You do not have to participate and you can refuse to answer any question. Even if you begin the web-based online survey, you can stop at any time.

**There are no foreseeable risks or discomforts to you for taking part in this study.**

**There are no direct benefits to you from participating in this study.** However, your responses may help us learn more about how state-of-the-art protein function prediction methods come across to on-the-field experts. It is hoped that this feedback shall be an invaluable resource to the community of computational biologists.

**As a token of our appreciation for completing the survey, you will receive a $10 Starbucks gift card by email after you have completed all 3 surveys.**

**Your part in this study will be handled in a confidential manner. No reports or publications based on this research will identify you or any individual as being affiliated with this project.**

**If you have any questions about this study,** please feel free to contact Rashika Ramola, the person mainly responsible for the research. You can also contact Prof. Predrag Radivojac, the Principal Investigator.

**If you have any questions regarding your rights as a research participant,** please contact Nan C. Regina, Director, Human Subject Research Protection, Mail Stop: 560-177, 360 Huntington Avenue, Northeastern University, Boston, MA 02115. Tel: 617.373.4588,. You may call anonymously if you wish.

**This study has been reviewed and approved by the Northeastern University Institutional Review Board (#19-10-08).**

**By checking the "I consent" button below you are indicating that you consent to participate in this study. Please print out a copy of this consent screen or download a copy of the consent form for your records.**

Thank you for your time.

Predrag Radivojac

### Appendix B - Experimentalists: General Form

This survey should take between 30 minutes and 1 hour. Please mark the time to help us see how long it took you.

Name (Optional)

Title (Optional)

Affiliated Institution(s): (Optional)

Note: The name, title and affiliated institution information will not be shared outside this research study even if you provide it.

Fields of specialization (check all that apply):

- Biology ▢
- Chemistry ▢
- Physics ▢
- Medicine ▢
- Mathematics ▢
- Statistics ▢
- Computer Science ▢
- Other ▢

1.2. Years of experience in your area of specialization:

O 0-2

O 2-5

O 5-10

O 10 or more

**X--------------------------------------------------------------------------------------------------X**

Here is a list of some databases used in bioinformatics.

Please indicate your level of familiarity with each one.
0 - not familiar; 1- heard of it, never used; 2- use rarely; 3- use sometimes; 4- use frequently

| | **0** | **1** | **2** | **3** | **4** |
|---|---|---|---|---|---|
| UniProt | O | O | O | O | O |
| Swiss-Prot | O | O | O | O | O |
| Gene Ontology | O | O | O | O | O |
| Brenda | O | O | O | O | O |
| DisProt | O | O | O | O | O |
| Protein Data Bank | O | O | O | O | O |
| Pfam | O | O | O | O | O |
| KEGG | O | O | O | O | O |
| Protein Data Bank | O | O | O | O | O |
| Ensembl | O | O | O | O | O |
| PATRIC | O | O | O | O | O |
| FlyBase | O | O | O | O | O |
| SGD | O | O | O | O | O |
| WormBase | O | O | O | O | O |
| BiGRID | O | O | O | O | O |

| | | | | | |
|---|---|---|---|---|---|
| SCOP | O | O | O | O | O |
| CATH | O | O | O | O | O |

What other bioinformatics databases or knowledge bases do you use?

Do you use the annotations of gene/protein function (such as Gene Ontology terms or Enzyme Commission numbers) in those databases for your research?

O Yes
O No

If yes to the previous question, do you consider the annotation's Evidence Codes when using those annotations in your research?

O Yes
O No
O What is Evidence Code?
O N/A

If yes to previous question, please answer the following two questions:

Have you ever used annotations with the "Inferred from Electronic Annotation (IEA)" evidence code in your research?

O Yes
O No
O N/A

What evidence codes do you trust the most?

If you are familiar with Gene Ontology (GO) please answer the following three questions. If not, skip to the next page.

How useful do you think is a GO annotation for an experimental scientist?

O 0 = not useful at all
O 1 = somewhat useful
O 2 = moderately useful
O 3 = very useful

How well do you think GO terms describe protein function?

O 0 = not well at all
O 1 = well enough
O 2 = very well

Do you have any further comments related to the previous two questions?

**X------------------------------------------------------------------------------------------------------X**

Familiarity with bioinformatics software

Here is a list of some software packages used in bioinformatics.

Please indicate your level of familiarity with each one.

0 - not familiar; 1- heard of it, never used; 2- use rarely; 3- use sometimes; 4- use frequently

| | | 0 | 1 | 2 | 3 | 4 |
|---|---|---|---|---|---|---|
| a. | BLAST | O | O | O | O | O |
| b. | DNAStar | O | O | O | O | O |
| c. | MEGA | O | O | O | O | O |
| d. | Clustal | O | O | O | O | O |
| e. | UCSC Genome Browser | O | O | O | O | O |

What bioinformatics software(s) do you use?

Briefly describe the purpose for which you use these software packages.

Do you use any software for the purpose of understanding a protein's function?

O Yes
O No

If yes, which software packages do you use:

### Appendix C - Experimentalists: Specific Form

How well do you know the following proteins on a scale from 0 to 5, where 0 = no knowledge at all and 5 = expert knowledge. Circle one number for each protein.

| | | 0 | 1 | 2 | 3 | 4 |
|---|---|---|---|---|---|---|
| a. | HBA1 (human) | O | O | O | O | O |
| b. | P53 (human) | O | O | O | O | O |
| c. | TPST1 (human) | O | O | O | O | O |
| d. | FOLR1 (human) | O | O | O | O | O |

There is a vast number of potential protein activities in and outside the cell. Gene Ontology (GO) terms standardize the description of protein functions at the molecular and biological level, in part to make the knowledge usable by computational methods. There are three sub-ontologies in GO: MFO (Molecular Function Ontology), BPO (Biological Process Ontology) and CCO (Cellular Component Ontology). Molecular function is the function at the molecular level (e.g. “catalytic activity” or “sodium channel activity”), biological process takes place at the level of pathways and biological processes (e.g. “apoptosis”, “glycolysis”), whereas cellular component describes where protein’s activity takes place (e.g. “nucleus”, “Golgi apparatus”).

In the next segment, you will be given ontological annotations for four proteins. Each has annotations in MFO, BPO and CCO together with confidence scores. We will ask you questions about these proteins.

The terms in GO are hierarchical. For example, “hydrolase activity” is a “catalytic activity” and therefore the two terms are connected by a relationship is-a in the graph. These relationships are visualized by arrows.

**Gene Ontology terms for hPNPase (MFO sub-ontology)**

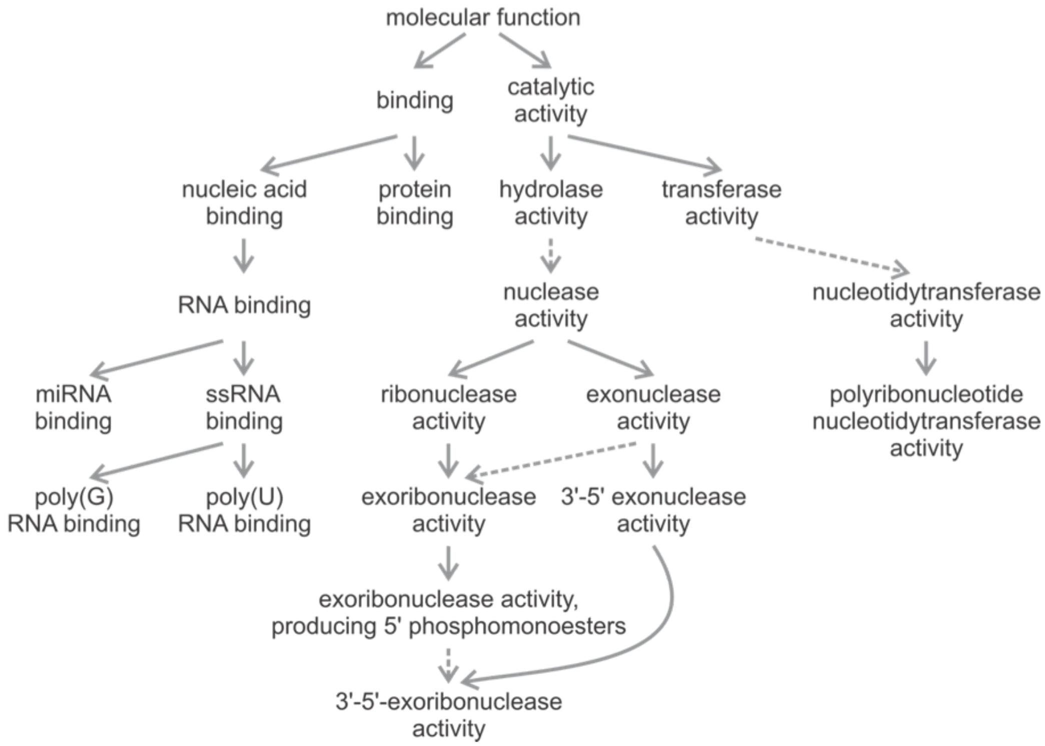

Figure modified from: Radivojac et al. Large-scale evaluation of protein function prediction. *Nat. Methods.* (2013) 10(3):221-227

X-----------------------------------------------------------------------------------------------X

You will now be given a set of Gene Ontology terms (Biological Processes, Molecular Functions and Cellular Components) for hemoglobin subunit alpha (HBA1) predicted by one protein function prediction algorithm. Each term will be preceded by a confidence score. High score indicates high confidence and low score indicates low confidence.

Please assign one or more of these categories to each Gene Ontology terms: Known (K), Useful (U), Surprising, possible (P), Surprising, doubtful (D), Wrong (W).

Please mark terms (using, K, U, P, D and/or W) as

a. Known (K): This is a well-known function of this protein
b. Useful (U): I find this prediction to be worthy of a follow-up study or it confirms my suspicion
c. Surprising, possible (P): I did not expect this prediction, but it's possible it is correct

d. Surprising, doubtful (D): I did not expect this prediction, but I very much doubt it is correct
e. Wrong (W): I believe this is a wrong prediction

#### Example:

#### Insulin-like growth factor 1 receptor (IGF1R) - Biological Process Ontology

| | Known (K) | Useful (U) | Surprising, possible (P) | Surprising, doubtful (D) | Wrong (W). |
|---|---|---|---|---|---|
| 0.91 glucose homeostasis | ✓ | ✓ | | | |
| 0.8 negative regulation of apoptotic process | | ✓ | ✓ | | |
| 0.70 immune system process | | | | | ✓ |

**HBA1 - Biological Process Ontology**

| | Known | Useful | Surprising, possible | Possible, doubtful | Wrong |
|---|---|---|---|---|---|
| 0.92 metabolic_process | ☐ | ☐ | ☐ | ☐ | ☐ |
| 0.91 transport | ☐ | ☐ | ☐ | ☐ | ☐ |
| 0.87 response_to_stimulus | ☐ | ☐ | ☐ | ☐ | ☐ |
| 0.83 cellular_response_to_stimulus | ☐ | ☐ | ☐ | ☐ | ☐ |
| 0.83 cell_communication | ☐ | ☐ | ☐ | ☐ | ☐ |
| 0.82 nitrogen_compound_metabolic_process | ☐ | ☐ | ☐ | ☐ | ☐ |
| 0.81 cellular_nitrogen_compound_metabolic_process | ☐ | ☐ | ☐ | ☐ | ☐ |
| 0.81 regulation_of_nitrogen_compound_metabolic_process | ☐ | ☐ | ☐ | ☐ | ☐ |
| 0.81 signaling | ☐ | ☐ | ☐ | ☐ | ☐ |
| 0.81 regulation_of_metabolic_process | ☐ | ☐ | ☐ | ☐ | ☐ |
| 0.8 establishment_of_localization_in_cell | ☐ | ☐ | ☐ | ☐ | ☐ |
| 0.8 cellular_metabolic_process | ☐ | ☐ | ☐ | ☐ | ☐ |
| 0.79 nucleobase-containing_compound_metabolic_process | ☐ | ☐ | ☐ | ☐ | ☐ |
| 0.78 ion_transport | ☐ | ☐ | ☐ | ☐ | ☐ |
| 0.77 developmental_process | ☐ | ☐ | ☐ | ☐ | ☐ |
| 0.77 biosynthetic_process | ☐ | ☐ | ☐ | ☐ | ☐ |
| 0.77 heterocycle_metabolic_process | ☐ | ☐ | ☐ | ☐ | ☐ |

| | | | | | |
|---|---|---|---|---|---|
| 0.76 regulation_of_RNA_metabolic_process | ☐ | ☐ | ☐ | ☐ | ☐ |
| 0.76 organelle_organization | ☐ | ☐ | ☐ | ☐ | ☐ |
| 0.76 regulation_of_RNA_biosynthetic_process | ☐ | ☐ | ☐ | ☐ | ☐ |
| 0.75 cellular_localization | ☐ | ☐ | ☐ | ☐ | ☐ |
| 0.75 regulation_of_nucleic_acid-templated_transcription | ☐ | ☐ | ☐ | ☐ | ☐ |
| 0.75 cellular_aromatic_compound_metabolic_process | ☐ | ☐ | ☐ | ☐ | ☐ |
| 0.75 anatomical_structure_development | ☐ | ☐ | ☐ | ☐ | ☐ |
| 0.75 establishment_of_protein_localization | ☐ | ☐ | ☐ | ☐ | ☐ |

**HBA1 - Cellular Component Ontology**

| | Known | Useful | Surprising, possible | Surprising, doubtful | Wrong |
|---|---|---|---|---|---|
| 0.96 intracellular_organelle | ☐ | ☐ | ☐ | ☐ | ☐ |
| 0.96 cytoplasm | ☐ | ☐ | ☐ | ☐ | ☐ |
| 0.91 intracellular_membrane-bounded_organelle | ☐ | ☐ | ☐ | ☐ | ☐ |
| 0.87 membrane | ☐ | ☐ | ☐ | ☐ | ☐ |
| 0.85 mitochondrial_membrane | ☐ | ☐ | ☐ | ☐ | ☐ |
| 0.84 mitochondrion | ☐ | ☐ | ☐ | ☐ | ☐ |
| 0.84 macromolecular_complex | ☐ | ☐ | ☐ | ☐ | ☐ |
| 0.81 nucleus | ☐ | ☐ | ☐ | ☐ | ☐ |
| 0.77 ribosome | ☐ | ☐ | ☐ | ☐ | ☐ |
| 0.76 protein_complex | ☐ | ☐ | ☐ | ☐ | ☐ |
| 0.74 cytosol | ☐ | ☐ | ☐ | ☐ | ☐ |
| 0.72 organelle_membrane | ☐ | ☐ | ☐ | ☐ | ☐ |
| 0.72 mitochondrial_envelope | ☐ | ☐ | ☐ | ☐ | ☐ |
| 0.7 nuclear_lumen | ☐ | ☐ | ☐ | ☐ | ☐ |
| 0.64 nucleoplasm | ☐ | ☐ | ☐ | ☐ | ☐ |
| 0.63 mitochondrial_inner_membrane | ☐ | ☐ | ☐ | ☐ | ☐ |
| 0.62 integral_component_of_membrane | ☐ | ☐ | ☐ | ☐ | ☐ |
| 0.57 extracellular_region | ☐ | ☐ | ☐ | ☐ | ☐ |
| 0.56 intrinsic_component_of_membrane | ☐ | ☐ | ☐ | ☐ | ☐ |
| 0.54 vesicle | ☐ | ☐ | ☐ | ☐ | ☐ |
| 0.51 membrane-bounded_vesicle | ☐ | ☐ | ☐ | ☐ | ☐ |

**HBA1 - Molecular Function Ontology**

| | Known | Useful | Surprising, possible | Surprising, doubtful | Wrong |
|---|---|---|---|---|---|
| 0.93 cytoskeletal_protein_binding | ☐ | ☐ | ☐ | ☐ | ☐ |
| 0.85 receptor_binding | ☐ | ☐ | ☐ | ☐ | ☐ |
| 0.85 protein_complex_binding | ☐ | ☐ | ☐ | ☐ | ☐ |
| 0.84 tubulin_binding | ☐ | ☐ | ☐ | ☐ | ☐ |
| 0.84 actin_binding | ☐ | ☐ | ☐ | ☐ | ☐ |
| 0.82 organic_cyclic_compound_binding | ☐ | ☐ | ☐ | ☐ | ☐ |
| 0.78 protein_domain_specific_binding | ☐ | ☐ | ☐ | ☐ | ☐ |
| 0.76 microtubule_binding | ☐ | ☐ | ☐ | ☐ | ☐ |
| 0.74 cation_binding | ☐ | ☐ | ☐ | ☐ | ☐ |
| 0.72 RNA_binding | ☐ | ☐ | ☐ | ☐ | ☐ |
| 0.68 transporter_activity | ☐ | ☐ | ☐ | ☐ | ☐ |
| 0.66 catalytic_activity | ☐ | ☐ | ☐ | ☐ | ☐ |
| 0.64 protein_heterodimerization_activity | ☐ | ☐ | ☐ | ☐ | ☐ |
| 0.63 cell_adhesion_molecule_binding | ☐ | ☐ | ☐ | ☐ | ☐ |
| 0.61 nucleoside-triphosphatase_activity | ☐ | ☐ | ☐ | ☐ | ☐ |
| 0.61 protein_kinase_binding | ☐ | ☐ | ☐ | ☐ | ☐ |
| 0.6 kinase_binding | ☐ | ☐ | ☐ | ☐ | ☐ |
| 0.59 nucleic_acid_binding | ☐ | ☐ | ☐ | ☐ | ☐ |
| 0.59 hydrolase_activity | ☐ | ☐ | ☐ | ☐ | ☐ |

Is any functional information about these proteins missing? Please provide it if you are aware of it.

X---------------------------------------------------------------------------------------------------------X

There are a vast number of potential protein activities in and outside the cell. Gene Ontology (GO) terms standardize the description of protein functions at the molecular and biological level, in part to make the knowledge usable by computational methods. There are three sub-ontologies in GO: MFO (Molecular Function Ontology), BPO (Biological Process Ontology) and CCO (Cellular Component Ontology). Molecular function is the function at the molecular level (e.g. “catalytic activity” or “sodium channel activity”), biological process takes place at the level of pathways and biological processes (e.g. “apoptosis”, “glycolysis”), whereas cellular component describes where protein’s activity takes place (e.g. “nucleus”, “Golgi apparatus”).

In the next segment, you will be given ontological annotations for four proteins. Each has annotations in MFO, BPO and CCO together with confidence scores. We will ask you questions about these proteins.

The terms in GO are hierarchical. For example, “hydrolase activity” is a “catalytic activity” and therefore the two terms are connected by a relationship is-a in the graph. These relationships are visualized by arrows.

##### Gene Ontology terms for hPNPase (MFO sub-ontology)

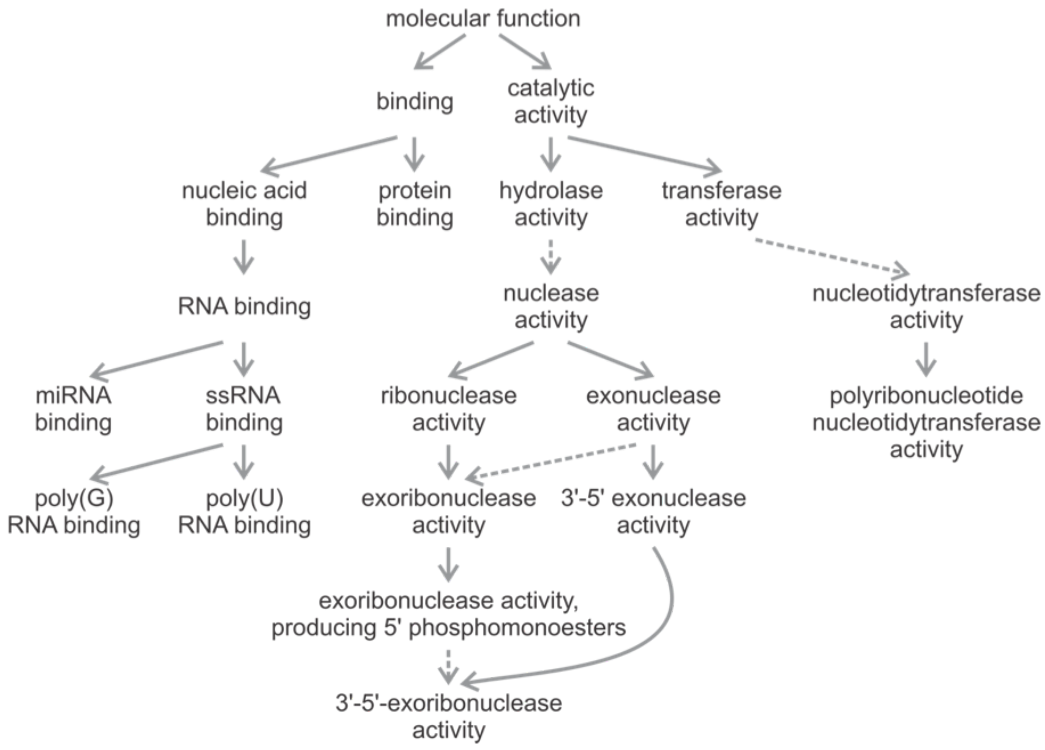

Figure modified from: Radivojac et al. Large-scale evaluation of protein function prediction. *Nat. Methods.* (2013) 10(3):221-227

X---------------------------------------------------------------------------------------------------X

You will now be given a set of Gene Ontology terms (Biological Processes, Molecular Functions and Cellular Components) for tumor protein (P53) predicted by one protein function prediction algorithm. Each term will be preceded by a confidence score. High score indicates high confidence and low score indicates low confidence.

Please assign one or more of these categories to each Gene Ontology terms: Known (K), Useful (U), Surprising, possible (P), Surprising, doubtful (D), Wrong (W).

Please mark terms (using, K, U, P, D and/or W) as

a. Known (K): This is a well-known function of this protein
b. Useful (U): I find this prediction to be worthy of a follow-up study or it confirms my suspicion
c. Surprising, possible (P): I did not expect this prediction, but it's possible it is correct
d. Surprising, doubtful (D): I did not expect this prediction, but I very much doubt it is correct

e. Wrong (W): I believe this is a wrong prediction

**Example:**

#### Insulin-like growth factor 1 receptor (IGF1R) - Biological Process Ontology

| | Known (K) | Useful (U) | Surprising, possible (P) | Surprising, doubtful (D) | Wrong (W). |
|---|---|---|---|---|---|
| 0.91 glucose homeostasis | ✓ | ✓ | | | |
| 0.8 negative regulation of apoptotic process | | ✓ | ✓ | | |
| 0.70 immune system process | | | | | ✓ |

#### P53 - Biological Process Ontology

| | Known | Useful | Surprising, possible | Surprising, doubtful | Wrong |
|---|---|---|---|---|---|
| 0.94 regulation_of_metabolic_process | ☐ | ☐ | ☐ | ☐ | ☐ |
| 0.93 cellular_metabolic_process | ☐ | ☐ | ☐ | ☐ | ☐ |
| 0.91 regulation_of_gene_expression | ☐ | ☐ | ☐ | ☐ | ☐ |
| 0.91 metabolic_process | ☐ | ☐ | ☐ | ☐ | ☐ |
| 0.88 cellular_nitrogen_compound_ metabolic_process | ☐ | ☐ | ☐ | ☐ | ☐ |
| 0.88 cellular_macromolecule_biosynthetic_process | ☐ | ☐ | ☐ | ☐ | ☐ |
| 0.87 RNA_splicing | ☐ | ☐ | ☐ | ☐ | ☐ |
| 0.86 regulation_of_RNA_biosynthetic_process | ☐ | ☐ | ☐ | ☐ | ☐ |
| 0.86 cell_communication | ☐ | ☐ | ☐ | ☐ | ☐ |
| 0.86 nitrogen_compound_metabolic_process | ☐ | ☐ | ☐ | ☐ | ☐ |
| 0.86 cellular_aromatic_compound_ metabolic_process | ☐ | ☐ | ☐ | ☐ | ☐ |
| 0.85 heterocycle_metabolic_process | ☐ | ☐ | ☐ | ☐ | ☐ |
| 0.85 gene_expression | ☐ | ☐ | ☐ | ☐ | ☐ |
| 0.85 nucleobase-containing_compound_metabolic_process | ☐ | ☐ | ☐ | ☐ | ☐ |
| 0.84 biosynthetic_process | ☐ | ☐ | ☐ | ☐ | ☐ |
| 0.83 regulation_of_nucleic_acid-templated_transcription | ☐ | ☐ | ☐ | ☐ | ☐ |
| 0.82 regulation_of_nitrogen_compound_ metabolic_process | ☐ | ☐ | ☐ | ☐ | ☐ |

| | | | | | |
|---|---|---|---|---|---|
| 0.81 response_to_stimulus | ☐ | ☐ | ☐ | ☐ | ☐ |
| 0.8 signal_transduction | ☐ | ☐ | ☐ | ☐ | ☐ |
| 0.8 signaling | ☐ | ☐ | ☐ | ☐ | ☐ |
| 0.8 cellular_response_to_stimulus | ☐ | ☐ | ☐ | ☐ | ☐ |
| 0.78 RNA_metabolic_process | ☐ | ☐ | ☐ | ☐ | ☐ |
| 0.77 developmental_process | ☐ | ☐ | ☐ | ☐ | ☐ |
| 0.77 macromolecule_biosynthetic_process | ☐ | ☐ | ☐ | ☐ | ☐ |
| 0.76 regulation_of_RNA_metabolic_process | ☐ | ☐ | ☐ | ☐ | ☐ |

**P53 - Cellular Component Ontology**

| | Known | Useful | Surprising, possible | Surprising, doubtful | Wrong |
|---|---|---|---|---|---|
| 0.94 intracellular_organelle | ☐ | ☐ | ☐ | ☐ | ☐ |
| 0.9 intracellular_membrane-bounded_ organelle | ☐ | ☐ | ☐ | ☐ | ☐ |
| 0.85 nuclear_lumen | ☐ | ☐ | ☐ | ☐ | ☐ |
| 0.85 nucleus | ☐ | ☐ | ☐ | ☐ | ☐ |
| 0.82 nucleolus | ☐ | ☐ | ☐ | ☐ | ☐ |
| 0.81 cytoplasm | ☐ | ☐ | ☐ | ☐ | ☐ |
| 0.78 nucleoplasm | ☐ | ☐ | ☐ | ☐ | ☐ |
| 0.76 macromolecular_complex | ☐ | ☐ | ☐ | ☐ | ☐ |
| 0.72 cytosol | ☐ | ☐ | ☐ | ☐ | ☐ |
| 0.69 protein_complex | ☐ | ☐ | ☐ | ☐ | ☐ |
| 0.57 spliceosomal_complex | ☐ | ☐ | ☐ | ☐ | ☐ |
| 0.55 nuclear_body | ☐ | ☐ | ☐ | ☐ | ☐ |

**P53 - Molecular Function Ontology**

| | Known | Useful | Surprising, possible | Surprising, doubtful | Wrong |
|---|---|---|---|---|---|
| 0.97 nucleic_acid_binding | ☐ | ☐ | ☐ | ☐ | ☐ |
| 0.95 organic_cyclic_compound_binding | ☐ | ☐ | ☐ | ☐ | ☐ |
| 0.92 nucleotide_binding | ☐ | ☐ | ☐ | ☐ | ☐ |
| 0.91 adenyl_nucleotide_binding | ☐ | ☐ | ☐ | ☐ | ☐ |
| 0.89 RNA_polymerase_II_transcription_regulatory_region_sequence-specific_DNA_binding_transcription_factor_activity_involved_in_positive_regulation_of_transcription | ☐ | ☐ | ☐ | ☐ | ☐ |
| 0.87 small_molecule_binding | ☐ | ☐ | ☐ | ☐ | ☐ |
| 0.87 DNA_binding | ☐ | ☐ | ☐ | ☐ | ☐ |
| 0.87 receptor_binding | ☐ | ☐ | ☐ | ☐ | ☐ |
| 0.8 metal_ion_binding | ☐ | ☐ | ☐ | ☐ | ☐ |
| 0.8 transition_metal_ion_binding | ☐ | ☐ | ☐ | ☐ | ☐ |
| 0.76 cation_binding | ☐ | ☐ | ☐ | ☐ | ☐ |
| 0.73 RNA_binding | ☐ | ☐ | ☐ | ☐ | ☐ |
| 0.69 sequence-specific_DNA_binding_transcription_factor_activity | ☐ | ☐ | ☐ | ☐ | ☐ |
| 0.68 kinase_binding | ☐ | ☐ | ☐ | ☐ | ☐ |
| 0.67 protein_complex_binding | ☐ | ☐ | ☐ | ☐ | ☐ |
| 0.67 sequence-specific_DNA_binding | ☐ | ☐ | ☐ | ☐ | ☐ |
| 0.66 poly(A)_RNA_binding | ☐ | ☐ | ☐ | ☐ | ☐ |

| | | | | | |
|---|---|---|---|---|---|
| 0.65 protein_heterodimerization_activity | ☐ | ☐ | ☐ | ☐ | ☐ |
| 0.64 nucleic_acid_binding_transcription_factor_activity | ☐ | ☐ | ☐ | ☐ | ☐ |
| 0.6 transcription_factor_binding | ☐ | ☐ | ☐ | ☐ | ☐ |
| 0.58 purine_nucleoside_binding | ☐ | ☐ | ☐ | ☐ | ☐ |
| 0.57 sequence-specific_DNA_binding_RNA_polymerase_II_transcription_factor_activity | ☐ | ☐ | ☐ | ☐ | ☐ |
| 0.56 cytoskeletal_protein_binding | ☐ | ☐ | ☐ | ☐ | ☐ |
| 0.55 RNA_polymerase_II_core_promoter_proximal_region_sequence-specific_DNA_binding_transcription_factor_activity_involved_in_positive_regulation_of_transcription | ☐ | ☐ | ☐ | ☐ | ☐ |
| 0.53 protein_kinase_binding | ☐ | ☐ | ☐ | ☐ | ☐ |

Is any functional information about these proteins missing? Please provide it if you are aware of it.

X-----------------------------------------------------------------------------------------------------------X

You will now be given a set of Gene Ontology terms (Biological Processes, Molecular Functions and Cellular Components) for Tyrosylprotein Sulfotransferase 1 (TPST1) predicted by one protein function prediction algorithm. Each term will be preceded by a confidence score. High score indicates high confidence and low score indicates low confidence.

Please assign one or more of these categories to each Gene Ontology terms: Known (K), Useful (U), Surprising, possible (P), Surprising, doubtful (D), Wrong (W).

Please mark terms (using, K, U, P, D and/or W) as

a. Known (K): This is a well-known function of this protein
b. Useful (U): I find this prediction to be worthy of a follow-up study or it confirms my suspicion
c. Surprising, possible (P): I did not expect this prediction, but it's possible it is correct
d. Surprising, doubtful (D): I did not expect this prediction, but I very much doubt it is correct
e. Wrong (W): I believe this is a wrong prediction

#### Example:

#### Insulin-like growth factor 1 receptor (IGF1R) - Biological Process Ontology

| | Known (K) | Useful (U) | Surprising, possible (P) | Surprising, doubtful (D) | Wrong (W). |
|---|---|---|---|---|---|
| 0.91 glucose homeostasis | ✓ | ✓ | | | |
| 0.8 negative regulation of apoptotic process | | ✓ | ✓ | | |
| 0.70 immune system process | | | | | ✓ |

**TPST1 - Biological Process Ontology**

| | Known | Useful | Surprising, possible | Surprising, doubtful | Wrong |
|---|---|---|---|---|---|
| 0.92 cellular_metabolic_process | ☐ | ☐ | ☐ | ☐ | ☐ |
| 0.91 response_to_stimulus | ☐ | ☐ | ☐ | ☐ | ☐ |
| 0.9 biosynthetic_process | ☐ | ☐ | ☐ | ☐ | ☐ |
| 0.89 metabolic_process | ☐ | ☐ | ☐ | ☐ | ☐ |
| 0.83 cellular_response_to_stimulus | ☐ | ☐ | ☐ | ☐ | ☐ |
| 0.81 signal_transduction | ☐ | ☐ | ☐ | ☐ | ☐ |
| 0.79 cellular_macromolecule_biosynthetic_process | ☐ | ☐ | ☐ | ☐ | ☐ |
| 0.79 developmental_process | ☐ | ☐ | ☐ | ☐ | ☐ |
| 0.77 signaling | ☐ | ☐ | ☐ | ☐ | ☐ |
| 0.76 small_molecule_metabolic_process | ☐ | ☐ | ☐ | ☐ | ☐ |
| 0.76 oxidation-reduction_process | ☐ | ☐ | ☐ | ☐ | ☐ |
| 0.72 regulation_of_metabolic_process | ☐ | ☐ | ☐ | ☐ | ☐ |
| 0.72 anatomical_structure_development | ☐ | ☐ | ☐ | ☐ | ☐ |
| 0.71 protein_metabolic_process | ☐ | ☐ | ☐ | ☐ | ☐ |
| 0.69 carboxylic_acid_metabolic_process | ☐ | ☐ | ☐ | ☐ | ☐ |
| 0.68 cell_communication | ☐ | ☐ | ☐ | ☐ | ☐ |
| 0.67 cellular_protein_metabolic_process | ☐ | ☐ | ☐ | ☐ | ☐ |
| 0.67 multicellular_organismal_development | ☐ | ☐ | ☐ | ☐ | ☐ |
| 0.66 nitrogen_compound_metabolic_process | ☐ | ☐ | ☐ | ☐ | ☐ |

| | | | | | |
|---|---|---|---|---|---|
| 0.65 transport | ☐ | ☐ | ☐ | ☐ | ☐ |
| 0.65 positive_regulation_of_metabolic_process | ☐ | ☐ | ☐ | ☐ | ☐ |
| 0.61 regulation_of_nitrogen_compound_metabolic_process | ☐ | ☐ | ☐ | ☐ | ☐ |
| 0.59 cell_differentiation | ☐ | ☐ | ☐ | ☐ | ☐ |
| 0.58 organic_acid_metabolic_process | ☐ | ☐ | ☐ | ☐ | ☐ |
| 0.58 cellular_protein_modification_process | ☐ | ☐ | ☐ | ☐ | ☐ |

**TPST1 - Cellular Component Ontology**

| | Known | Useful | Surprising, possible | Surprising, doubtful | Wrong |
|---|---|---|---|---|---|
| 0.97 membrane | ☐ | ☐ | ☐ | ☐ | ☐ |
| 0.96 integral_component_of_membrane | ☐ | ☐ | ☐ | ☐ | ☐ |
| 0.96 intracellular_organelle | ☐ | ☐ | ☐ | ☐ | ☐ |
| 0.93 intracellular_membrane-bounded_organelle | ☐ | ☐ | ☐ | ☐ | ☐ |
| 0.92 cytoplasm | ☐ | ☐ | ☐ | ☐ | ☐ |
| 0.88 organelle_membrane | ☐ | ☐ | ☐ | ☐ | ☐ |
| 0.87 Golgi_membrane | ☐ | ☐ | ☐ | ☐ | ☐ |
| 0.85 intrinsic_component_of_membrane | ☐ | ☐ | ☐ | ☐ | ☐ |
| 0.79 mitochondrial_envelope | ☐ | ☐ | ☐ | ☐ | ☐ |
| 0.79 mitochondrial_membrane | ☐ | ☐ | ☐ | ☐ | ☐ |
| 0.78 extracellular_region | ☐ | ☐ | ☐ | ☐ | ☐ |
| 0.77 endoplasmic_reticulum_membrane | ☐ | ☐ | ☐ | ☐ | ☐ |
| 0.76 mitochondrion | ☐ | ☐ | ☐ | ☐ | ☐ |
| 0.74 endomembrane_system | ☐ | ☐ | ☐ | ☐ | ☐ |
| 0.74 bounding_membrane_of_organelle | ☐ | ☐ | ☐ | ☐ | ☐ |
| 0.74 nuclear_outer_membrane-endoplasmic_reticulum_membrane_network | ☐ | ☐ | ☐ | ☐ | ☐ |
| 0.73 mitochondrial_inner_membrane | ☐ | ☐ | ☐ | ☐ | ☐ |
| 0.66 endoplasmic_reticulum | ☐ | ☐ | ☐ | ☐ | ☐ |

| | | | | | |
|---|---|---|---|---|---|
| 0.62 vesicle | ☐ | ☐ | ☐ | ☐ | ☐ |
| 0.58 plasma_membrane | ☐ | ☐ | ☐ | ☐ | ☐ |
| 0.55 membrane-bounded_vesicle | ☐ | ☐ | ☐ | ☐ | ☐ |
| 0.55 Golgi_apparatus | ☐ | ☐ | ☐ | ☐ | ☐ |
| 0.52 protein_complex | ☐ | ☐ | ☐ | ☐ | ☐ |
| 0.51 extracellular_vesicular_exosome | ☐ | ☐ | ☐ | ☐ | ☐ |

**TPST1 - Molecular Function Ontology**

| | Known | Useful | Surprising, possible | Surprising, doubtful | Wrong |
|---|---|---|---|---|---|
| 0.96 transferase_activity_transferring_hexosyl_groups | ☐ | ☐ | ☐ | ☐ | ☐ |
| 0.94 catalytic_activity | ☐ | ☐ | ☐ | ☐ | ☐ |
| 0.93 metal_ion_binding | ☐ | ☐ | ☐ | ☐ | ☐ |
| 0.92 iron_ion_binding | ☐ | ☐ | ☐ | ☐ | ☐ |
| 0.89 transferase_activity | ☐ | ☐ | ☐ | ☐ | ☐ |
| 0.89 organic_cyclic_compound_binding | ☐ | ☐ | ☐ | ☐ | ☐ |
| 0.86 oxidoreductase_activity | ☐ | ☐ | ☐ | ☐ | ☐ |
| 0.78 cation_binding | ☐ | ☐ | ☐ | ☐ | ☐ |
| 0.77 adenyl_nucleotide_binding | ☐ | ☐ | ☐ | ☐ | ☐ |
| 0.74 heme_binding | ☐ | ☐ | ☐ | ☐ | ☐ |
| 0.71 transferase_activity_transferring_glycosyl_groups | ☐ | ☐ | ☐ | ☐ | ☐ |
| 0.71 nucleoside_binding | ☐ | ☐ | ☐ | ☐ | ☐ |
| 0.68 purine_nucleoside_binding | ☐ | ☐ | ☐ | ☐ | ☐ |
| 0.67 receptor_binding | ☐ | ☐ | ☐ | ☐ | ☐ |
| 0.67 small_molecule_binding | ☐ | ☐ | ☐ | ☐ | ☐ |
| 0.65 oxidoreductase_activity_acting_on_paired_donors | ☐ | ☐ | ☐ | ☐ | ☐ |
| 0.64 ribonucleoside_binding | ☐ | ☐ | ☐ | ☐ | ☐ |

| | | | | | |
|---|---|---|---|---|---|
| 0.64 nucleotide_binding | ☐ | ☐ | ☐ | ☐ | ☐ |
| 0.60 protein_complex_binding | ☐ | ☐ | ☐ | ☐ | ☐ |
| 0.60 ATP_binding | ☐ | ☐ | ☐ | ☐ | ☐ |
| 0.59 hydrolase_activity | ☐ | ☐ | ☐ | ☐ | ☐ |
| 0.55 purine_nucleotide_binding | ☐ | ☐ | ☐ | ☐ | ☐ |
| 0.51 purine_ribonucleoside_triphosphate_binding | ☐ | ☐ | ☐ | ☐ | ☐ |

Is any functional information about these proteins missing? Please provide it if you are aware of it.

X-----------------------------------------------------------------------------------------------------------X

You will now be given a set of Gene Ontology terms (Biological Processes, Molecular Functions and Cellular Components) for folate receptor 1 (FOLR1) predicted by one protein function prediction algorithm. Each term will be preceded by a confidence score. High score indicates high confidence and low score indicates low confidence.

Please assign one or more of these categories to each Gene Ontology terms: Known (K), Useful (U), Surprising, possible (P), Surprising, doubtful (D), Wrong (W).

Please mark terms (using, K, U, P, D and/or W) as

a. Known (K): This is a well-known function of this protein
b. Useful (U): I find this prediction to be worthy of a follow-up study or it confirms my suspicion
c. Surprising, possible (P): I did not expect this prediction, but it's possible it is correct

d. Surprising, doubtful (D): I did not expect this prediction, but I very much doubt it is correct
e. Wrong (W): I believe this is a wrong prediction

##### Example:

##### Insulin-like growth factor 1 receptor (IGF1R) - Biological Process Ontology

| | Known (K) | Useful (U) | Surprising, possible (P) | Surprising, doubtful (D) | Wrong (W). |
|---|---|---|---|---|---|
| 0.91 glucose homeostasis | ✓ | ✓ | | | |
| 0.8 negative regulation of apoptotic process | | ✓ | ✓ | | |
| 0.70 immune system process | | | | | ✓ |

**FOLR1 - Biological Process Ontology**

| | Known | Useful | Surprising, possible | Surprising, doubtful | Wrong |
|---|---|---|---|---|---|
| 0.96 transferase_activity_transferring_hexosyl_groups | ☐ | ☐ | ☐ | ☐ | ☐ |
| 0.94 catalytic_activity | ☐ | ☐ | ☐ | ☐ | ☐ |
| 0.93 metal_ion_binding | ☐ | ☐ | ☐ | ☐ | ☐ |
| 0.92 iron_ion_binding | ☐ | ☐ | ☐ | ☐ | ☐ |
| 0.89 transferase_activity | ☐ | ☐ | ☐ | ☐ | ☐ |
| 0.89 organic_cyclic_compound_binding | ☐ | ☐ | ☐ | ☐ | ☐ |
| 0.86 oxidoreductase_activity | ☐ | ☐ | ☐ | ☐ | ☐ |
| 0.78 cation_binding | ☐ | ☐ | ☐ | ☐ | ☐ |
| 0.77 adenyl_nucleotide_binding | ☐ | ☐ | ☐ | ☐ | ☐ |
| 0.74 heme_binding | ☐ | ☐ | ☐ | ☐ | ☐ |
| 0.71 transferase_activity_transferring_glycosyl_groups | ☐ | ☐ | ☐ | ☐ | ☐ |
| 0.71 nucleoside_binding | ☐ | ☐ | ☐ | ☐ | ☐ |
| 0.68 purine_nucleoside_binding | ☐ | ☐ | ☐ | ☐ | ☐ |
| 0.67 receptor_binding | ☐ | ☐ | ☐ | ☐ | ☐ |
| 0.67 small_molecule_binding | ☐ | ☐ | ☐ | ☐ | ☐ |
| 0.65 oxidoreductase_activity_acting_on_paired_donors | ☐ | ☐ | ☐ | ☐ | ☐ |

| | | | | | |
|---|---|---|---|---|---|
| 0.64 ribonucleoside_binding | ☐ | ☐ | ☐ | ☐ | ☐ |
| 0.64 nucleotide_binding | ☐ | ☐ | ☐ | ☐ | ☐ |
| 0.60 protein_complex_binding | ☐ | ☐ | ☐ | ☐ | ☐ |
| 0.60 ATP_binding | ☐ | ☐ | ☐ | ☐ | ☐ |
| 0.59 hydrolase_activity | ☐ | ☐ | ☐ | ☐ | ☐ |
| 0.55 purine_nucleotide_binding | ☐ | ☐ | ☐ | ☐ | ☐ |
| 0.51 purine_ribonucleoside_triphosphate_binding | ☐ | ☐ | ☐ | ☐ | ☐ |

**FOLR1 - Cellular Component Ontology**

| | Known | Useful | Surprising, possible | Surprising, doubtful | Wrong |
|---|---|---|---|---|---|
| 0.97 intrinsic_component_of_membrane | ☐ | ☐ | ☐ | ☐ | ☐ |
| 0.95 membrane | ☐ | ☐ | ☐ | ☐ | ☐ |
| 0.92 integral_component_of_membrane | ☐ | ☐ | ☐ | ☐ | ☐ |
| 0.91 extracellular_region | ☐ | ☐ | ☐ | ☐ | ☐ |
| 0.89 plasma_membrane | ☐ | ☐ | ☐ | ☐ | ☐ |
| 0.88 cell_periphery | ☐ | ☐ | ☐ | ☐ | ☐ |
| 0.85 cytoplasm | ☐ | ☐ | ☐ | ☐ | ☐ |
| 0.85 vesicle | ☐ | ☐ | ☐ | ☐ | ☐ |
| 0.84 extracellular_vesicular_exosome | ☐ | ☐ | ☐ | ☐ | ☐ |
| 0.83 intracellular_membrane-bounded_organelle | ☐ | ☐ | ☐ | ☐ | ☐ |
| 0.81 organelle_membrane | ☐ | ☐ | ☐ | ☐ | ☐ |
| 0.80 integral_component_of_plasma_membrane | ☐ | ☐ | ☐ | ☐ | ☐ |
| 0.80 intracellular_organelle | ☐ | ☐ | ☐ | ☐ | ☐ |
| 0.76 endomembrane_system | ☐ | ☐ | ☐ | ☐ | ☐ |
| 0.75 extracellular_space | ☐ | ☐ | ☐ | ☐ | ☐ |
| 0.75 membrane-bounded_vesicle | ☐ | ☐ | ☐ | ☐ | ☐ |
| 0.74 intrinsic_component_of_plasma_membrane | ☐ | ☐ | ☐ | ☐ | ☐ |
| 0.73 external_side_of_plasma_membrane | ☐ | ☐ | ☐ | ☐ | ☐ |

| | | | | | |
|---|---|---|---|---|---|
| 0.66 endoplasmic_reticulum | ☐ | ☐ | ☐ | ☐ | ☐ |
| 0.56 Golgi_membrane | ☐ | ☐ | ☐ | ☐ | ☐ |
| 0.53 bounding_membrane_of_organelle | ☐ | ☐ | ☐ | ☐ | ☐ |
| 0.51 cell_surface | ☐ | ☐ | ☐ | ☐ | ☐ |
| 0.51 cytoplasmic_vesicle | ☐ | ☐ | ☐ | ☐ | ☐ |

#### FOLR1 - Molecular Function Ontology

| | Known | Useful | Surprising, possible | Surprising, doubtful | Wrong |
|---|---|---|---|---|---|
| 0.95 metal_ion_binding | ☐ | ☐ | ☐ | ☐ | ☐ |
| 0.95 zinc_ion_binding | ☐ | ☐ | ☐ | ☐ | ☐ |
| 0.94 catalytic_activity | ☐ | ☐ | ☐ | ☐ | ☐ |
| 0.89 growth_factor_activity | ☐ | ☐ | ☐ | ☐ | ☐ |
| 0.89 G-protein_coupled_receptor_binding | ☐ | ☐ | ☐ | ☐ | ☐ |
| 0.85 receptor_binding | ☐ | ☐ | ☐ | ☐ | ☐ |
| 0.83 cytokine_activity | ☐ | ☐ | ☐ | ☐ | ☐ |
| 0.82 transferase_activity_transferring_glycosyl_groups | ☐ | ☐ | ☐ | ☐ | ☐ |
| 0.81 receptor_activity | ☐ | ☐ | ☐ | ☐ | ☐ |
| 0.80 cation_binding | ☐ | ☐ | ☐ | ☐ | ☐ |
| 0.77 organic_cyclic_compound_binding | ☐ | ☐ | ☐ | ☐ | ☐ |
| 0.75 glycosaminoglycan_binding | ☐ | ☐ | ☐ | ☐ | ☐ |
| 0.74 hydrolase_activity | ☐ | ☐ | ☐ | ☐ | ☐ |
| 0.74 transferase_activity | ☐ | ☐ | ☐ | ☐ | ☐ |
| 0.71 small_molecule_binding | ☐ | ☐ | ☐ | ☐ | ☐ |
| 0.70 transferase_activity_transferring_glycosyl_groups | ☐ | ☐ | ☐ | ☐ | ☐ |
| 0.69 transporter_activity | ☐ | ☐ | ☐ | ☐ | ☐ |
| 0.65 protein_complex_binding | ☐ | ☐ | ☐ | ☐ | ☐ |
| 0.62 transition_metal_ion_binding | ☐ | ☐ | ☐ | ☐ | ☐ |

| | | | | | |
|---|---|---|---|---|---|
| 0.62 enzyme_regulator_activity | ☐ | ☐ | ☐ | ☐ | ☐ |
| 0.57 hydrolase_activity_acting_on_ester_ bondsactivity | ☐ | ☐ | ☐ | ☐ | ☐ |

Is any functional information about these proteins missing? Please provide it if you are aware of it.

X---------------------------------------------------------------------------------------------------------------X

How would you describe these predictions?

Do you think the algorithm has done a good job?

Do you think the scores given to terms by the software were reasonable?

Would you prefer to see these predictions in a graphical format, that also provides visualization of the ontology? (see Figure 1)

When investigating the function of a particular protein, what do you do? Describe databases you use and generally the steps you take. If you use any algorithms, tools, or web site, please mention those as well.

If there was an ideal algorithm, how should it communicate protein function to you?

### Appendix D - Experimentalists: Comparative Form

In this segment, you will see up to 25 Gene Ontology (GO) predictions for TPST1 and FOLR1 from two algorithms. Algorithm 1 is the same one you saw in the previous section. Algorithm 2 is different. After showing these predictions, we will ask you a few questions to compare the two algorithms.

**Protein**: TPST1 in human
**Ontology**: Biological Process Ontology (BPO)

| Algorithm 1 | | Algorithm 2 | |
|---|---|---|---|
| 0.92 | cellular_metabolic_process | 1 | biological_process |
| 0.91 | response_to_stimulus | 0.88 | cellular process |
| 0.9 | biosynthetic_process | 0.75 | metabolic process |
| 0.89 | metabolic_process | 0.71 | cellular metabolic process |
| 0.83 | cellular_response_to_stimulus | 0.7 | organic substance metabolic process |
| 0.81 | signal_transduction | 0.62 | primary metabolic process |
| 0.79 | cellular_macromolecule_ biosynthetic_process | 0.59 | nitrogen compound metabolic process |
| 0.79 | developmental_process | 0.46 | biosynthetic process |
| 0.77 | signaling | 0.46 | organonitrogen compound metabolic process |
| 0.76 | small_molecule_metabolic_ process | 0.46 | organic substance biosynthetic process |
| 0.76 | oxidation-reduction_process | 0.45 | cellular nitrogen compound metabolic process |
| 0.72 | regulation_of_metabolic_process | 0.45 | cellular biosynthetic process |
| 0.72 | anatomical_structure_development | 0.4 | macromolecule metabolic process |

| | | | |
|---|---|---|---|
| 0.71 | protein_metabolic_process | 0.35 | organonitrogen compound biosynthetic process |
| 0.69 | carboxylic_acid_metabolic_ process | 0.34 | cellular macromolecule metabolic process |
| 0.68 | cell_communication | 0.34 | organic cyclic compound metabolic process |
| 0.67 | cellular_protein_metabolic_ process | 0.32 | cellular aromatic compound metabolic process |
| 0.67 | multicellular_organismal_ development | 0.32 | heterocycle metabolic process |
| 0.66 | nitrogen_compound_ metabolic_process | 0.3 | cellular nitrogen compound biosynthetic process |
| 0.65 | transport | 0.29 | small molecule metabolic process |
| 0.65 | positive_regulation_of_ metabolic_process | 0.29 | single-organism metabolic process |
| 0.61 | regulation_of_nitrogen_ compound_metabolic_process | 0.26 | nucleobase-containing compound metabolic process |
| 0.59 | cell_differentiation | 0.24 | gene expression |
| 0.58 | organic_acid_metabolic_process | 0.23 | macromolecule biosynthetic process |
| 0.58 | cellular_protein_ modification_process | 0.23 | protein metabolic process |

How would you rate Algorithm 1 and Algorithm 2 based on the quality of  predictions, quality of assigned scores and the completeness of predictions? The ratings are on a scale of 1 to five, where 1 = terrible, 2 = unsatisfactory, 3 = good, 4 = very good, 5 = excellent.

| | Quality of predictions on a scale of 1 to 5 | | | | | Qualtiy of allotted scores on a scale of 1 to 5 | | | | | Completeness of predictions on a scale of 1 to 5 | | | | |
|---|---|---|---|---|---|---|---|---|---|---|---|---|---|---|---|
| | 1 | 2 | 3 | 4 | 5 | 1 | 2 | 3 | 4 | 5 | 1 | 2 | 3 | 4 | 5 |
| Algorithm 1 | ○ | ○ | ○ | ○ | ○ | ○ | ○ | ○ | ○ | ○ | ○ | ○ | ○ | ○ | ○ |
| Algorithm 2 | ○ | ○ | ○ | ○ | ○ | ○ | ○ | ○ | ○ | ○ | ○ | ○ | ○ | ○ | ○ |

**Protein**: TPST1 in human
**Ontology**: Cellular Component Ontology (CCO)

| Algorithm 1 | | Algorithm 2 | |
|---|---|---|---|
| 0.97 | membrane | 1 | cellular_component |
| 0.96 | integral_component_of_membrane | 0.75 | intracellular |
| 0.96 | intracellular_organelle | 0.75 | cell |
| 0.93 | intracellular_membrane-bounded_organelle | 0.75 | cell part |
| 0.92 | cytoplasm | 0.56 | cytoplasm |
| 0.88 | organelle_membrane | 0.56 | intracellular part |
| 0.87 | Golgi_membrane | 0.39 | organelle |
| 0.85 | intrinsic_component_of_membrane | 0.38 | intracellular organelle |
| 0.79 | mitochondrial_envelope | 0.3 | membrane |
| 0.79 | mitochondrial_membrane | 0.26 | membrane-bounded organelle |
| 0.78 | extracellular_region | 0.24 | intracellular membrane-bounded organelle |
| 0.77 | endoplasmic_reticulum_membrane | 0.2 | integral to membrane |
| 0.76 | mitochondrion | 0.2 | intrinsic to membrane |
| 0.74 | endomembrane_system | 0.2 | macromolecular complex |
| 0.74 | bounding_membrane_of_organelle | 0.2 | membrane part |

| | | | |
|---|---|---|---|
| 0.74 | nuclear_outer_membrane-endoplasmic_reticulum_membrane_network | 0.19 | non-membrane-bounded organelle |
| 0.73 | mitochondrial_inner_membrane | 0.19 | intracellular non-membrane-bounded organelle |
| 0.66 | endoplasmic_reticulum | 0.19 | cell periphery |
| 0.62 | vesicle | 0.18 | plasma membrane |
| 0.58 | plasma_membrane | 0.12 | nucleus |
| 0.55 | membrane-bounded_vesicle | 0.12 | ribosome |
| 0.55 | Golgi_apparatus | 0.12 | ribonucleoprotein complex |
| 0.52 | protein_complex | 0.12 | cytoplasmic part |
| 0.51 | extracellular_vesicular_exosome | 0.09 | extracellular region |

How would you rate Algorithm 1 and Algorithm 2 based on the quality of predictions, quality of assigned scores and the completeness of predictions? The ratings are on a scale of 1 to five, where 1 = terrible, 2 = unsatisfactory, 3 = good, 4 = very good, 5 = excellent.

| | Quality of predictions on a scale of 1 to 5 | | | | | Qualtiy of allotted scores on a scale of 1 to 5 | | | | | Completeness of predictions on a scale of 1 to 5 | | | | |
|---|---|---|---|---|---|---|---|---|---|---|---|---|---|---|---|
| | 1 | 2 | 3 | 4 | 5 | 1 | 2 | 3 | 4 | 5 | 1 | 2 | 3 | 4 | 5 |
| Algorithm 1 | ○ | ○ | ○ | ○ | ○ | ○ | ○ | ○ | ○ | ○ | ○ | ○ | ○ | ○ | ○ |
| Algorithm 2 | ○ | ○ | ○ | ○ | ○ | ○ | ○ | ○ | ○ | ○ | ○ | ○ | ○ | ○ | ○ |

**Protein**: TPST1 in human
**Ontology**: Molecular Function Ontology (MFO)

| Algorithm 1 | | Algorithm 2 | |
|---|---|---|---|
| 0.96 | transferase_activity_transferring_hexosyl_groups | 1 | molecular_function |
| 0.94 | catalytic_activity | 0.7 | binding |
| 0.93 | metal_ion_binding | 0.64 | catalytic activity |
| 0.92 | iron_ion_binding | 0.49 | organic cyclic compound binding |
| 0.89 | transferase_activity | 0.49 | heterocyclic compound binding |
| 0.89 | organic_cyclic_compound_binding | 0.45 | ion binding |
| 0.86 | oxidoreductase_activity | 0.28 | small molecule binding |
| 0.78 | cation_binding | 0.27 | cation binding |
| 0.77 | adenyl_nucleotide_binding | 0.26 | nucleotide binding |
| 0.74 | heme_binding | 0.26 | nucleic acid binding |
| 0.71 | transferase_activity_transferring_glycosyl_groups | 0.26 | anion binding |
| 0.71 | nucleoside_binding | 0.26 | metal ion binding |
| 0.68 | purine_nucleoside_binding | 0.26 | nucleoside phosphate binding |
| 0.67 | receptor_binding | 0.24 | transferase activity |

| | | | |
|---|---|---|---|
| 0.67 | small_molecule_binding | 0.23 | carbohydrate derivative binding |
| 0.65 | oxidoreductase_activity_acting_on_paired_donors_with_incorporation_or_reduction_of_molecular_oxygen | 0.22 | purine nucleotide binding |
| 0.64 | ribonucleoside_binding | 0.22 | ribonucleotide binding |
| 0.64 | nucleotide_binding | 0.22 | purine ribonucleotide binding |
| 0.6 | protein_complex_binding | 0.21 | purine ribonucleoside triphosphate binding |
| 0.6 | ATP_binding | 0.19 | nucleoside binding |
| 0.59 | hydrolase_activity | 0.19 | purine nucleoside binding |
| 0.55 | purine_nucleotide_binding | 0.19 | ATP binding |
| 0.51 | purine_ribonucleoside_triphosphate_binding | 0.19 | hydrolase activity |

How would you rate Algorithm 1 and Algorithm 2 based on the quality of predictions, quality of assigned scores and the completeness of predictions? The ratings are on a scale of 1 to five, where 1 = terrible, 2 = unsatisfactory, 3 = good, 4 = very good, 5 = excellent.

| | Quality of predictions on a scale of 1 to 5 | | | | | Qualtiy of allotted scores on a scale of 1 to 5 | | | | | Completeness of predictions on a scale of 1 to 5 | | | | |
|---|---|---|---|---|---|---|---|---|---|---|---|---|---|---|---|
| | 1 | 2 | 3 | 4 | 5 | 1 | 2 | 3 | 4 | 5 | 1 | 2 | 3 | 4 | 5 |
| Algorithm 1 | ○ | ○ | ○ | ○ | ○ | ○ | ○ | ○ | ○ | ○ | ○ | ○ | ○ | ○ | ○ |
| Algorithm 2 | ○ | ○ | ○ | ○ | ○ | ○ | ○ | ○ | ○ | ○ | ○ | ○ | ○ | ○ | ○ |

**Protein**: FOLR1 in human
**Ontology**: Biological Process Ontology (BPO)

| Algorithm 1 | | Algorithm 2 | |
|---|---|---|---|
| 0.94 | response_to_stimulus | 1 | biological_process |
| 0.84 | signal_transduction | 0.88 | cellular process |
| 0.84 | cellular_response_to_stimulus | 0.75 | metabolic process |
| 0.81 | developmental_process | 0.71 | cellular metabolic process |
| 0.81 | metabolic_process | 0.7 | organic substance metabolic process |
| 0.79 | transport | 0.62 | primary metabolic process |
| 0.78 | nitrogen_compound_metabolic_process | 0.59 | nitrogen compound metabolic process |
| 0.78 | anatomical_structure_development | 0.46 | biosynthetic process |
| 0.76 | regulation_of_metabolic_process | 0.46 | organonitrogen compound metabolic process |
| 0.76 | multicellular_organismal_development | 0.46 | organic substance biosynthetic process |
| 0.75 | signaling | 0.45 | cellular nitrogen compound metabolic process |
| 0.75 | cellular_metabolic_process | 0.45 | cellular biosynthetic process |
| 0.73 | cell_surface_receptor_signaling_pathway | 0.4 | macromolecule metabolic process |

**Protein**: FOLR1 in human
**Ontology**: Cellular Component Ontology (CCO)

| | | | |
|---|---|---|---|
| 0.75 | extracellular_space | 0.2 | membrane part |
| 0.75 | membrane-bounded_vesicle | 0.19 | non-membrane-bounded organelle |
| 0.74 | intrinsic_component_of_plasma _membrane | 0.19 | intracellular non-membrane-bounded organelle |
| 0.73 | external_side_of_plasma_ membrane | 0.19 | cell periphery |
| 0.66 | endoplasmic_reticulum | 0.18 | plasma membrane |
| 0.56 | Golgi_membrane | 0.12 | nucleus |
| 0.53 | bounding_membrane_of_organelle | 0.12 | ribosome |
| 0.51 | cell_surface | 0.12 | ribonucleoprotein complex |
| 0.51 | cytoplasmic_vesicle | 0.12 | cytoplasmic part |

How would you rate Algorithm 1 and Algorithm 2 based on the quality of predictions, quality of assigned scores and the completeness of predictions? The ratings are on a scale of 1 to five, where 1 = terrible, 2 = unsatisfactory, 3 = good, 4 = very good, 5 = excellent.

| | Quality of predictions on a scale of 1 to 5 | | | | | Qualtiy of allotted scores on a scale of 1 to 5 | | | | | Completeness of predictions on a scale of 1 to 5 | | | | |
|---|---|---|---|---|---|---|---|---|---|---|---|---|---|---|---|
| | 1 | 2 | 3 | 4 | 5 | 1 | 2 | 3 | 4 | 5 | 1 | 2 | 3 | 4 | 5 |
| Algorithm 1 | ○ | ○ | ○ | ○ | ○ | ○ | ○ | ○ | ○ | ○ | ○ | ○ | ○ | ○ | ○ |
| Algorithm 2 | ○ | ○ | ○ | ○ | ○ | ○ | ○ | ○ | ○ | ○ | ○ | ○ | ○ | ○ | ○ |

**Protein**: FOLR1 in human
**Ontology**: Molecular Function Ontology (MFO)

| | Algorithm 1 | | Algorithm 2 |
|---|---|---|---|
| 0.95 | metal_ion_binding | 1 | molecular_function |
| 0.95 | zinc_ion_binding | 0.7 | binding |
| 0.94 | catalytic_activity | 0.64 | catalytic activity |
| 0.89 | growth_factor_activity | 0.49 | organic cyclic compound binding |
| 0.89 | G-protein_coupled_ receptor_binding | 0.49 | heterocyclic compound binding |
| 0.85 | receptor_binding | 0.45 | ion binding |
| 0.83 | cytokine_activity | 0.28 | small molecule binding |
| 0.82 | transferase_activity_ transferring _glycosyl_groups | 0.27 | cation binding |
| 0.81 | receptor_activity | 0.26 | nucleotide binding |
| 0.8 | cation_binding | 0.26 | nucleic acid binding |
| 0.77 | organic_cyclic_compound_ binding | 0.26 | anion binding |
| 0.75 | glycosaminoglycan_binding | 0.26 | metal ion binding |
| 0.74 | hydrolase_activity | 0.26 | nucleoside phosphate binding |
| 0.74 | transferase_activity | 0.24 | transferase activity |

| | | | |
|---|---|---|---|
| 0.71 | small_molecule_binding | 0.23 | carbohydrate derivative binding |
| 0.7 | transferase_activity_ transferring _hexosyl_groups | 0.22 | purine nucleotide binding |
| 0.69 | transporter_activity | 0.22 | ribonucleotide binding |
| 0.65 | protein_complex_binding | 0.22 | purine ribonucleotide binding |
| 0.62 | transition_metal_ion_binding | 0.21 | purine ribonucleoside triphosphate binding |
| 0.62 | enzyme_regulator_activity | 0.19 | nucleoside binding |
| 0.57 | hydrolase_activity_acting_on_ ester_bonds | 0.19 | purine nucleoside binding |

How would you rate Algorithm 1 and Algorithm 2 based on the quality of predictions, quality of assigned scores and the completeness of predictions? The ratings are on a scale of 1 to five, where 1 = terrible, 2 = unsatisfactory, 3 = good, 4 = very good, 5 = excellent.

| | Quality of predictions on a scale of 1 to 5 | | | | | Qualtiy of allotted scores on a scale of 1 to 5 | | | | | Completeness of predictions on a scale of 1 to 5 | | | | |
|---|---|---|---|---|---|---|---|---|---|---|---|---|---|---|---|
| | 1 | 2 | 3 | 4 | 5 | 1 | 2 | 3 | 4 | 5 | 1 | 2 | 3 | 4 | 5 |
| Algorithm 1 | ○ | ○ | ○ | ○ | ○ | ○ | ○ | ○ | ○ | ○ | ○ | ○ | ○ | ○ | ○ |
| Algorithm 2 | ○ | ○ | ○ | ○ | ○ | ○ | ○ | ○ | ○ | ○ | ○ | ○ | ○ | ○ | ○ |

Do you have any other comments about performance of the two algorithms when compared to each other?

### Appendix E - Computational Biologists: Consent Form

**Northeastern University, Khoury College of Computer Sciences**

**Name of Investigator(s):** Prof. Predrag Radivojac

**Title of Project:** Assessing the usability and value of protein function prediction algorithms

**Sponsor:** National Science Foundation

**Information Sheet**

**Request to Participate in Research**

We would like to invite you to participate in a web-based online survey. The survey is part of a research study whose purpose is to understand the perception and utility of computational protein function prediction methods.

The surveys should take about 15 minutes to complete.

We are asking you to participate in this study because you are a computational biologist with experience in building bioinformatics tools. **You must be at least 18 years old to take this survey.**

**The decision to participate in this research project is voluntary.** You do not have to participate and you can refuse to answer any question. Even if you begin the web-based online survey, you can stop at any time.

**There are no foreseeable risks or discomforts to you for taking part in this study.**

**There are no direct benefits to you from participating in this study.** However, your responses may help us learn more about the approach of computational biologists towards the development of tools for protein function prediction. It is hoped that this feedback shall be an invaluable resource to the community of computational biologists.

**As a token of our appreciation for completing the survey, you will receive a $10 Starbucks gift card by email after you have completed all 3 surveys.**

**Your part in this study will be handled in a confidential manner. No reports or publications based on this research will identify you or any individual as being affiliated with this project.**

**If you have any questions about this study,** please feel free to contact Rashika Ramola, the person mainly responsible for the research. You can also contact Prof. Predrag Radivojac, the Principal Investigator.

**If you have any questions regarding your rights as a research participant,** please contact Nan C. Regina, Director, Human Subject Research Protection, Mail Stop: 560-177, 360 Huntington Avenue, Northeastern University, Boston, MA 02115. Tel: 617.373.4588,. You may call anonymously if you wish.

**This study has been reviewed and approved by the Northeastern University Institutional Review Board (#19-10-08).**

**By checking the "I consent" button below you are indicating that you consent to participate in this study. Please print out a copy of this consent screen or download a copy of the consent form for your records.**

Thank you for your time.

Predrag Radivojac

### Appendix F - Computational Biologists: General Form

This survey should take no more than 30 minutes. Please mark the time to help us see how long it took you.

Name (Optional)

Title (Optional)

Affiliated Institution(s): (Optional)

Note: The name, title and affiliated institution information will not be shared outside this research study even if you provide it.

Fields of specialization (check all that apply):

| | |
|---|---|
| Biology | ▢ |
| Chemistry | ▢ |
| Physics | ▢ |
| Medicine | ▢ |
| Mathematics | ▢ |
| Statistics | ▢ |
| Computer Science | ▢ |
| Other | ▢ |

1.2. Years of experience in your area of specialization:

O 0-2

O 2-5

O 5-10

O 10 or more

**X-----------------------------------------------------------------------------------------------X**
Here is a list of some databases used in bioinformatics.

Please indicate your level of familiarity with each one.
0 - not familiar; 1- heard of it, never used; 2- use rarely; 3- use sometimes; 4- use frequently

| | **0** | **1** | **2** | **3** | **4** |
|---|---|---|---|---|---|
| UniProt | O | O | O | O | O |
| Swiss-Prot | O | O | O | O | O |
| Gene Ontology | O | O | O | O | O |
| Brenda | O | O | O | O | O |
| DisProt | O | O | O | O | O |
| Protein Data Bank | O | O | O | O | O |
| Pfam | O | O | O | O | O |
| KEGG | O | O | O | O | O |
| Protein Data Bank | O | O | O | O | O |
| Ensembl | O | O | O | O | O |
| PATRIC | O | O | O | O | O |
| FlyBase | O | O | O | O | O |
| SGD | O | O | O | O | O |

| | | | | | |
|---|---|---|---|---|---|
| WormBase | O | O | O | O | O |
| BiGRID | O | O | O | O | O |
| SCOP | O | O | O | O | O |
| CATH | O | O | O | O | O |

What other bioinformatics databases or knowledge bases do you use?

Do you use the annotations of gene/protein function (such as Gene Ontology terms or Enzyme Commission classification numbers) in those databases for your research?

O Yes
O No

If yes to the previous question, do you consider the annotation’s Evidence Codes when using those annotations in your research?

O Yes
O No
O What is an Evidence Code?
O N/A

If yes to previous question, please answer the following two questions:

Have you ever used annotations with the “Inferred from Electronic Annotation (IEA)” evidence code in your research?

O Yes
O No
O N/A

What evidence codes do you trust the most?

If you are familiar with Gene Ontology (GO) please answer the following three questions. If not, skip to the next page.

How useful do you think is a GO annotation for an experimental scientist?

O 0 = not useful at all
O 1 = somewhat useful
O 2 = moderately useful
O 3 = very useful

How well do you think GO terms describe protein function?

O 0 = not well at all
O 1 = well enough
O 2 = very well

Do you have any further comments related to the previous two questions?

**X-----------------------------------------------------------------------------------------------------------X**

Familiarity with bioinformatics software

Below is a list of some software packages used in bioinformatics. Please indicate your level of familiarity with each one.

0 - not familiar; 1- heard of it, never used; 2- use rarely; 3- use sometimes; 4- use frequently

| | | 0 | 1 | 2 | 3 | 4 |
|---|---|---|---|---|---|---|
| a. | BLAST | O | O | O | O | O |
| b. | DNAStar | O | O | O | O | O |
| c. | MEGA | O | O | O | O | O |
| d. | Clustal | O | O | O | O | O |
| e. | UCSC Genome Browser | O | O | O | O | O |

Which bioinformatics software(s) do you use?

Briefly describe the purpose for which you use these software packages.

Do you use any software for the purpose of understanding a protein's function?

O Yes
O No

If yes, which software packages do you use:

**X-------------------------------------------------------------------------------------X**

What types of bioinformatics software does your lab develop?

Who do you think are the users of these bioinformatics tools?

Please assess your level of interaction with experimental scientists when you write software

O 0 = Not at all
O 1 = Minor
O 2 = Occasional
O 3 = Extensive

If you interacted with the users of the tools you develop, what feedback have you received from them? Have you incorporated the feedback and if not, why?

Do you think that developing tools for protein function prediction is an important problem?

O 0 = No
O 1 = It is somewhat important
O 2 = It is quite important
O 3 = It is key to understanding and driving biology

If the previous summary was not descriptive, please provide any additional thoughts

Does your lab develop protein function prediction algorithms?

If you answered yes to the previous question, what do you consider the distinctive feature of your algorithm?

How do you think the results of a protein function prediction pipeline should be presented? How is your tool presenting them?

When investigating a specific protein, what do you consider should be the typical steps that an experimental scientist should follow?

X-----------------------------------------------------------------------X

To what extent do you think CAFA (Critical Assessment of Function Annotations) is useful?

O 0 - never heard of CAFA
O 1- no useful
O 2- somewhat useful
O 3- highly useful

How good are the evaluation metrics used in CAFA for protein function prediction?

O 0 = They do not capture anything relevant
O 1 = They capture some relevant information
O 2 = They capture enough information to be relevant for some purposes
O 3 = They capture most relevant information.

Would you like to see more metrics for evaluating function prediction methods and if so, what should they reflect?

What do you think are the chief bottlenecks in protein function prediction? Check all that apply:

O Quality of data
O Ontologies
O Methodology
O Evaluation
O Other (elaborate, add a text field)

Anything else you would like to add?

### Appendix G - Biocurators: Consent Form

**Northeastern University, Khoury College of Computer Sciences**

**Name of Investigator(s):** Prof. Predrag Radivojac

**Title of Project:** Assessing the usability and value of protein function prediction algorithms

**Sponsor:** National Science Foundation

**Information Sheet**

**Request to Participate in Research**

We would like to invite you to participate in a web-based online survey. The survey is part of a research study whose purpose is to understand the perception and utility of computational protein function prediction methods.

The surveys should take about 15 minutes to complete.

We are asking you to participate in this study because of your work in biocuration. **You must be at least 18 years old to take this survey.**

**The decision to participate in this research project is voluntary.** You do not have to participate and you can refuse to answer any question. Even if you begin the web-based online survey, you can stop at any time.

**There are no foreseeable risks or discomforts to you for taking part in this study.**

**There are no direct benefits to you from participating in this study.** However, your responses may help us learn more about the approach of biocuators towards curating gene ontologies. It is hoped that this feedback shall be an invaluable resource to the community of computational biologists.

**As a token of our appreciation for completing the survey, you will receive a $10 Starbucks gift card by email after you have completed all 3 surveys.**

**Your part in this study will be handled in a confidential manner. No reports or publications based on this research will identify you or any individual as being affiliated with this project.**

**If you have any questions about this study,** please feel free to contact Rashika Ramola, the person mainly responsible for the research. You can also contact Prof. Predrag Radivojac, the Principal Investigator.

**If you have any questions regarding your rights as a research participant,** please contact Nan C. Regina, Director, Human Subject Research Protection, Mail Stop: 560-177, 360 Huntington Avenue, Northeastern University, Boston, MA 02115. Tel: 617.373.4588,. You may call anonymously if you wish.

**This study has been reviewed and approved by the Northeastern University Institutional Review Board (#19-10-08).**

**By checking the "I consent" button below you are indicating that you consent to participate in this study. Please print out a copy of this consent screen or download a copy of the consent form for your records.**

Thank you for your time.

Predrag Radivojac

### Appendix H - Biocurators: General Form

This survey should take no more than 30 minutes. Please mark the time to help us see how long it took you.

Name (Optional)

Title (Optional)

Affiliated Institution(s): (Optional)

Note: The name, title and affiliated institution information will not be shared outside this research study even if you provide it.

Fields of specialization (check all that apply):

| | |
|---|---|
| Biology | ▢ |
| Chemistry | ▢ |
| Physics | ▢ |
| Medicine | ▢ |
| Mathematics | ▢ |
| Statistics | ▢ |
| Computer Science | ▢ |
| Other | ▢ |

1.2. Years of experience in your area of specialization:

O 0-2

O 2-5

O 5-10

O 10 or more

**X---------------------------------------------------------------------------------------------------X**

Here is a list of some databases used in bioinformatics.

Please indicate your level of familiarity with each one.
0 - not familiar; 1- heard of it, never used; 2- use rarely; 3- use sometimes; 4- use frequently

| | **0** | **1** | **2** | **3** | **4** |
|---|---|---|---|---|---|
| UniProt | O | O | O | O | O |
| Swiss-Prot | O | O | O | O | O |
| Gene Ontology | O | O | O | O | O |
| Brenda | O | O | O | O | O |
| DisProt | O | O | O | O | O |
| Protein Data Bank | O | O | O | O | O |
| Pfam | O | O | O | O | O |
| KEGG | O | O | O | O | O |
| Protein Data Bank | O | O | O | O | O |
| Ensembl | O | O | O | O | O |
| PATRIC | O | O | O | O | O |
| FlyBase | O | O | O | O | O |
| SGD | O | O | O | O | O |
| WormBase | O | O | O | O | O |

| | | | | | |
|---|---|---|---|---|---|
| BiGRID | O | O | O | O | O |
| SCOP | O | O | O | O | O |
| CATH | O | O | O | O | O |

What other bioinformatics databases or knowledge bases do you use?

How reliable do you think are database annotations labeled with “Inferred from Electronic Annotation (IEA)” evidence code?

O 0 = completely unreliable;
O 1 = somewhat reliable;
O 2 = pretty reliable;
O 3 = almost as good as experimental annotations

What evidence codes do you trust the most?

If you are familiar with Gene Ontology (GO) please answer the following three questions. If not, skip to the next page.

How useful do you think is a GO annotation for an experimental scientist?

O 0 = not useful at all
O 1 = somewhat useful
O 2 = moderately useful
O 3 = very useful

How useful do you think is a GO annotation for a computational scientist?

O 0 = not useful at all
O 1 = somewhat useful
O 2 = moderately useful
O 3 = very useful

How well do you think GO terms describe protein function?

O 0 = not well at all
O 1 = well enough
O 2 = very well

Do you have any further comments related to the previous three questions?

**X----------------------------------------------------------------------------------------------------------X**

Familiarity with bioinformatics software

Below is a list of some software packages used in bioinformatics. Please indicate your level of familiarity with each one.

0 - not familiar; 1- heard of it, never used; 2- use rarely; 3- use sometimes; 4- use frequently

| | | 0 | 1 | 2 | 3 | 4 |
|---|---|---|---|---|---|---|
| a. | BLAST | O | O | O | O | O |
| b. | DNAStar | O | O | O | O | O |
| c. | MEGA | O | O | O | O | O |
| d. | Clustal | O | O | O | O | O |

| | | | | | | |
|---|---|---|---|---|---|---|
| e. | UCSC Genome Browser | O | O | O | O | O |

Which bioinformatics software(s) do you use?

Briefly describe the purpose for which you use these software packages.

Do you use any software for the purpose of understanding a protein's function?

O Yes
O No

If yes, which software packages do you use:

**X------------------------------------------------------------------------------------X**

Please assess your level of interaction with experimental scientists when you curate ontologies:

O 0 = not at all
O 1 = minor
O 2 = occasional
O 3 = extensive

Please assess your level of interaction with computational scientists when you curate ontologies:

O 0 = Not at all
O 1 = Minor
O 2 = Occasional
O 3 = Extensive

If you interacted with the users of the ontologies and databases you develop, what feedback have you received from them? Have you incorporated it and if not, why?

Do you think that developing tools for protein function prediction is an important problem?

O 0 = No; It is idiosyncratic
O 1 = It is somewhat important
O 2 = It is quite important
O 3 = It holds one of the keys to understanding and driving biology

If the previous summary was not descriptive, please provide any additional thoughts:

Does your lab use protein function prediction algorithms?

If yes to the previous question, what algorithm(s) do you find reliable?

How do you think the results of a protein function pipeline should be presented?

To what extent do you think experimental scientists use ontologies and databases the way you envision?

O 0 = Not familiar with use
O 1 = They often use it inappropriately
O 2 = They use it somewhat appropriately
O 3 = They use it very appropriately

How good are evaluation metrics used in CAFA for protein function prediction?
O 0 = Do not capture anything relevant
O 1 = What is CAFA?
O 2 = Capture some relevant information
O 3 = Capture enough information for some good decision making
O 4 = Capture most relevant information

Would you like to see more metrics for function prediction and if so, what should they reflect?

What do you think is the bottleneck in protein function prediction? Check all that apply:

O Quality of data

O Ontologies
O Methodology
O Evaluation

Do you have any further comments on the previous question
